# Using human genetics to develop strategies to increase erythropoietic output from genome-edited hematopoietic stem and progenitor cells

**DOI:** 10.1101/2023.08.04.552064

**Authors:** Sofia E. Luna, Joab Camarena, Jessica P. Hampton, Kiran R. Majeti, Carsten T. Charlesworth, Eric Soupene, Sridhar Selvaraj, Kun Jia, Vivien A. Sheehan, M. Kyle Cromer, Matthew H. Porteus

**Author notes:** Contributed equally (Sofia E. Luna, Joab Camarena).

## Abstract

Human genetic polymorphisms result in a diversity of phenotypes. Some sequences are pathologic and lead to monogenic diseases, while others may confer beneficial traits. Genome editing is a powerful tool to recreate genotypes found in the population, including the ability to correct pathologic mutations. One of the best characterized naturally occurring mutations causing congenital erythrocytosis arises from a truncation in the erythropoietin receptor (tEPOR) which can result in non-pathogenic hyper-production of red blood cells (RBCs). Using the precision of CRISPR/Cas9 genome editing, we have recreated tEPOR and studied the effect of variations of the genotype on RBC development. We then combined tEPOR with a correction strategy developed for β-thalassemia and demonstrated that coupling the two genome editing events gave RBCs a significant selective advantage. This demonstrates the potential of combining human genetics with the precision of genome editing to enable safer and more effective genome editing therapies for patients with serious genetic diseases.

## Main

Many of the initial applications of clinical genome editing have aimed to correct or compensate for disease-causing mutations of monogenic diseases. However, human genetic variation is more nuanced than monogenic diseases, as there are variants that appear to confer positive health benefits as well. Examples are people that have bi-allelic deletions in *CCR5* causing resistance to HIV infection, a variety of polymorphisms causing upregulation of fetal hemoglobin, and mutations in *PCSK9* causing low cholesterol levels^1–3^. It is important, however, to broadly assess variations that occur in only small numbers of people for both their potential risks as well as for their potential benefits.

Congenital erythrocytosis (CE) is a rare phenotype in which people have higher than normal levels of red blood cells and consequently elevated hemoglobin. While there are multiple genetic variants that can lead to this condition, perhaps the best characterized genotype was first identified in the family of a Finnish Olympic gold medal-winning cross-country skier who was found to have levels of hemoglobin >50% higher than normal^4^. This was attributed to truncations in the erythropoietin receptor (tEPOR) that eliminate elements of the intracellular inhibitory domain to erythropoietin (EPO) signaling^4,5^. This domain contains binding sites for SHP1 which normally leads to downregulation of EPO-dependent JAK2/STAT5 signaling (Extended Data Figure 1a). Further studies have shown that tEPOR does not create a constitutively active EPOR signaling cascade, but rather imparts hypersensitivity to EPO^5,6^. As a consequence, these kindreds with tEPOR typically present with abnormally low levels of EPO, indicating a new homeostasis is attained to prevent CE from becoming pathogenic. There have been reports of thrombotic and hemorrhagic events likely due to erythrocytosis but many of these have a benign clinical course^7^. More importantly, families with CE have not shown an increased predisposition to cancer, demonstrating that this is not a pre-malignant genetic condition^8^.

While prior studies, including those by the Smithies group and Tisdale group, have investigated the effects of viral-mediated delivery and expression of *tEPOR*^9–11^, random insertion into the genomes of billions of hematopoietic stem and progenitor cells (HSPCs) in the context of bone marrow transplant presents a serious safety concern, and has resulted in a “Black Box” warning in the United States for lovotibeglogene autotemcel, a lentiviral gene therapy drug approved for sickle cell disease^12^. In addition, such instances of viral-mediated delivery require expression of *tEPOR* using a non-native, exogenous promoter, which departs from native *EPOR* regulation and has the potential for unintended consequences such as pathogenic polycythemia. Nonetheless, viral-mediated expression studies provide a proof-of-concept that demonstrates that *tEPOR* expression can lend a selective advantage to transduced cells and provide a foundation for the utilization of more advanced genome engineering modalities.

Genome editing is a powerful method that enables the precise changing of nucleotides in the DNA of a cell. There are multiple genome-editing strategies including nuclease-based insertion/deletion (indel) formation, base editing, and prime editing, but the most versatile approach to genome-editing is homology-directed repair (HDR). In HDR, a nuclease-induced double-strand break (DSB) is repaired using a donor template. The natural donor template for HDR is the sister chromatid and the natural repair pathway is homologous recombination. By providing a donor template that resembles a sister chromatid with large homology arms flanking the intended cut site, the homologous recombination machinery can use this “substitute” sister chromatid to repair the DSB. HDR editing is the most flexible because it can create single nucleotide changes, precisely insert large gene cassettes, and even swap out large genomic regions for other gene sequences^13–16^. We utilize all of these applications of HDR in this work, including direct creation of the naturally occurring variant found in a human kindred.

One of the challenges in hematopoietic stem cell gene therapy is to achieve sufficient engraftment of the genetically engineered cells to have a beneficial clinical effect without increasing risk. To make this possible, effort is exerted to maximize editing frequencies in HSPCs^17–19^. Even if clinically relevant editing frequencies are achieved, high-morbidity chemo-therapeutic regimens are currently required to create niche space in the bone marrow (BM) for these edited HSPCs, which can create toxicities, including oncogenic risk^20–22^. In this work, we aimed to give edited cells a selective advantage such that low levels of engraftment might still result in a clinical benefit, perhaps enabling less toxic conditioning, through use of genome editing to recreate the CE phenotype by engineering tEPOR into HSPCs in different ways. We find that when tEPOR is engineered into human HSPCs using genome-editing, there is a significant selective advantage to the derived red blood cells (RBCs). We then show that this selective advantage can be coupled to a therapeutic gene edit to give the cells with the therapeutic edit a selective advantage in RBC development, without affecting the stem and progenitor cells. In this way, we demonstrate the power of combining human genetics with precision genome editing to potentially enable safer and more effective genome editing therapies for patients with serious genetic diseases, particularly those of red blood cells.

## Results

### Cas9-guided EPOR truncation in HSPCs enhances erythroid proliferation

Truncating mutations in the *EPOR* gene that cause clinically benign congenital erythrocytosis^4,8^ provide a potentially safe avenue to increase erythropoietic output from genome-edited HSPCs. In this study we designed Cas9 single-guide RNAs (sgRNAs)^17^ (termed *EPOR*-sg1 and *EPOR*-sg2) that overlap the location of the originally identified nonsense mutation, *EPOR* c.1316G>A (p.W439X) (“Mäntyranta variant”)^4^. (Figure 1a, Extended Data Figure 2a). Our hypothesis was that targeting this site in exon 8 with Cas9 would create a spectrum of indels, a subset of which would result in a frameshift of the reading frame and yield premature downstream stop codons in the *EPOR* gene.

**Figure 1:**
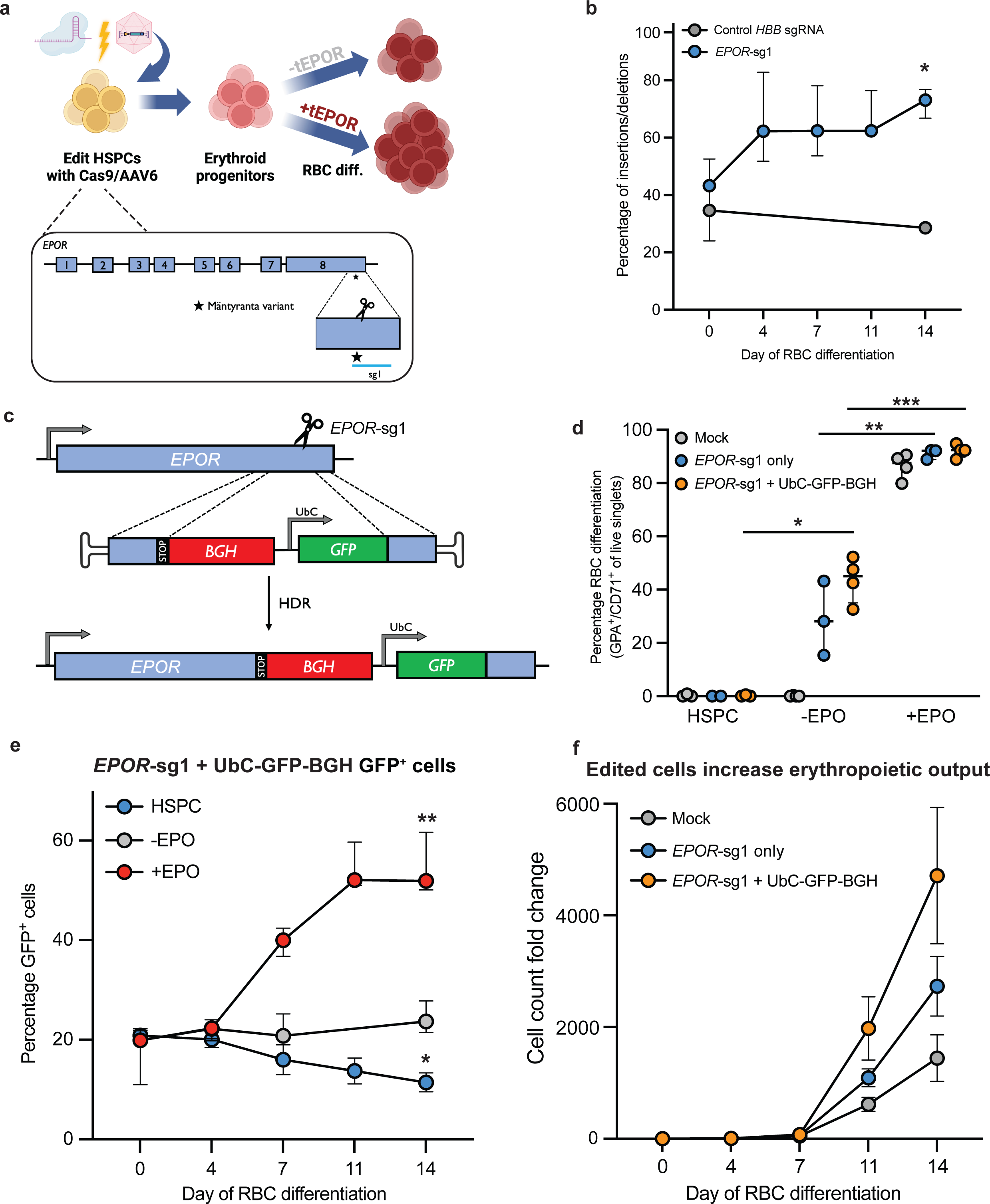
Cas9-guided *EPOR* truncation in HSPCs enhances erythroid proliferation. **a.** Schematic of HSPC editing and model of tEPOR effect. Representation of *EPOR* gene and location of the candidate sgRNA (*EPOR*-sg1) indicated by a line. Location of c.1316G>A mutation is denoted by the star. **b.** Frequency of indels created by sg1 in primary human CD34^+^ HSPCs over the course of erythroid differentiation compared to control *HBB* sgRNA. Points represent median ± interquartile range. Values represent biologically independent HSPC donors: N=5 for *EPOR*-sg1, N=1 for control *HBB* sgRNA. *P=0.0016 of day 1 vs. day 14 by unpaired two-tailed *t*-test. **c.** Genome editing strategy when using an AAV6 DNA repair template to introduce the *EPOR* c.1316G>A mutation followed by a BGH-polyA region and UbC-driven GFP reporter. **d.** Percentage of GPA^+^/CD71^+^ of live single cells on day 14 of differentiation. Bars represent median ± interquartile range. Values represent biologically independent HSPC donors: N=2-3 for HSPC, N=3-4 for -EPO and +EPO conditions. *P=0.0016 of -EPO vs. HSPC conditions; **P=0.003 and ***P=0.0001 of -EPO vs. +EPO conditions by unpaired two-tailed *t*-test. **e.** Percentage of GFP^+^ cells of live single cells maintained in RBC media +/- EPO or HSPC media as determined by flow cytometry. Points represent median ± interquartile range. Values represent biologically independent HSPC donors N=2 for HSPC condition and N=3-4 for -EPO and +EPO conditions. *P=0.04, **P=0.0006 of day 0 vs. day 14 by unpaired two-tailed *t*-test. **f.** Fold change in cell count throughout RBC differentiation (e.g., if at day 0 starting cell numbers were 1E5 cells total, therefore a fold count change of 1000 would yield a total cell number of 1E8 at day 14). Points represent mean ± SEM. Values represent biologically independent HSPC donors N=3 for Mock and *EPOR*-sg1 + BGH and N=2 for *EPOR*-sg1.

To test this hypothesis, we pre-complexed each sgRNA with high-fidelity Cas9 protein^23^ and delivered these ribonucleoprotein (RNP) complexes to human CD34^+^ HSPCs. At 2-3 days post-editing, we transferred cells into culture media that promotes erythroid differentiation over the course of two weeks^24^ (Figure 1a). To determine whether edited HSPCs have a proliferative advantage compared to unedited cells, we harvested genomic DNA at days 0, 4, 7, 11, and 14 of RBC differentiation. We then quantified indel frequency by PCR amplification followed by Sanger sequencing and decomposition analysis using TIDE^25^. In the absence of a selective advantage or disadvantage, percent edited alleles in cells at the beginning and end of RBC differentiation will be roughly equivalent—which is what we observe for the editing frequency of the *HBB* sgRNA used for correction of sickle cell disease. However, we observe that the editing frequency of *EPOR*-targeting sgRNAs increases significantly over the course of erythroid differentiation, to a greater extent in *EPOR*-sg1 than *EPOR*-sg2 (Figure 1b, Extended Data Figure 2b) (P=0.0016 for *EPOR*-sg1 from day 0 to day 14). Additionally, we show that the increase in indels for both *EPOR*-sg1 and *EPOR*-sg2 are predominantly driven by indels that yield downstream stop codons (Extended Data Figure 2c-d, Extended Data Figure 3a-b). This indicates that edited cells, particularly those with premature stop codons, are outcompeting unedited cells because of the EPO hypersensitivity of tEPOR-expressing cells in culture^5,6^.

Because not all indels created by the sgRNAs cause truncations in *EPOR*, we speculated that we could increase this proliferative effect by using HDR to insert a stop codon at the exact location of the original variant (c.1316G>A). To accomplish this, we designed an AAV6 repair template vector that introduces a stop codon into *EPOR* at the 439th amino acid (W439X) followed by a BGH-polyA tail to terminate transcription. We also included a downstream GFP marker driven by the constitutive human UbC promoter to ensure that each GFP^+^ allele harbors the intended *EPOR*-truncating mutation. The entire integration cassette was flanked by 950-base pair homology arms that corresponded to the genomic DNA immediately upstream and downstream of the intended Cas9 cut site created by the more effective *EPOR*-sg1 (Figure 1c). To determine whether this editing strategy was also able to drive enrichment of genome-edited RBCs, we complexed *EPOR*-sg1 with Cas9 protein and delivered this by electroporation to human CD34^+^ HSPCs followed by transduction with AAV6 DNA repair template. Two to three days post-editing we either maintained cells in HSPC media or began erythroid differentiation with 3 U/mL of EPO (+EPO), as has been previously described^14^, or with 0 U/mL of EPO (-EPO) in order to determine whether *tEPOR*-expressing cells retain EPO sensitivity or became EPO independent during their differentiation. At day 14 of erythroid differentiation, we stained for established RBC markers^14^ and analyzed cells using flow cytometry. We observed no differentiation when cells were kept in HSPC media and efficient RBC differentiation in all treatments with EPO. In the edited conditions in the absence of EPO, we observed moderate differentiation which may indicate the hypersensitivity of tEPOR-expressing cells to trace amounts of EPO in the media, as has been previously observed^4^ (Figure 1d). By analyzing GFP^+^ cells over the course of RBC differentiation, in the *EPOR*^W439X^-edited conditions we observed a significant increase in the frequency of edited cells in only the +EPO conditions (P=0.0006 comparing day 0 to day 14) (Figure 1e). At the end of differentiation, by quantifying both GFP^+^ cells and frequency of indel formation, we estimate that almost all the RBCs are derived from cells with a truncated EPOR due to either an indel or a UbC-GFP knock-in event (Extended Data Figure 4a). In addition to the competitive advantage that *tEPOR* expression gives to edited cells over the course of RBC differentiation, we also observed increased RBC production in both *EPOR*-sg1 and *EPOR*-sg1 + UbC-GFP-BGH conditions compared to Mock control in the +EPO condition (average 1.45x10^3^ total fold increase in mock edited cells versus 4.71x10^3^ in *EPOR*-sg1 + UbC-GFP-BGH over the 14-day RBC differentiation) (Figure 1f). These increased cell counts were not observed in the -EPO condition or when cells were maintained in HSPC media, indicating an EPO-driven increase in erythroid proliferation in cells expressing *tEPOR* (Extended Data Figure 4b). We also assessed whether a gradient of EPO concentrations (0 U/mL – 20 U/mL) over the course of RBC differentiation yielded varying degrees of enrichment of edited cells (Extended Data Figure 5a). On day 14, when compared to 0 U/mL EPO, we found minimal differences in GFP^+^ cells in the 1, 3, and 20 U/mL conditions and only a minor reduction in GFP^+^ cells in the 0.3 U/mL EPO condition (Extended Data Figure 5b-c), indicating that even low levels of EPO is sufficient to impart a selective advantage to edited cells consistent with prior work on the natural variants^5,6^. We also observed comparable levels of RBC differentiation at all concentrations except 0 U/mL EPO (Extended Data Figure 5d).

In terms of the safety of this editing strategy, we show that introduction of these *EPOR* indels yields RBCs with production of both fetal and adult hemoglobin tetramers following hemoglobin tetramer high-performance liquid chromatography (HPLC) (Extended Data Figure 6a). We also found similar colony number and lineage distribution from CD34^+^ HSPCs plated into wells containing methylcellulose media either with or without EPO that were scored for colony-formation ability after 14 days (Extended Data Figure 6b). As expected, there was a marked decrease in the ability to form BFU-E colonies in the absence of EPO even if the cells contained the tEPOR. These data reinforce the idea that truncation of EPOR does not alter the lineage bias of HSPCs, but rather has an effect only after commitment to the erythroid lineage. Additionally, although transient delivery of high-fidelity Cas9 has been shown to be highly specific to the on-target site^26^, we also evaluated potential off-target effects of the *EPOR*-sg1-RNP complex in HSPCs. We found that 94% (73/78) of candidate off-target sites with scores previously shown to be most informative for identifying sites with potential off-target activity^27^ resided in intergenic or intronic regions of the genome (Extended Data Figure 7a-b). We further interrogated potential off-target activity at the five sites that resided in exonic or UTR regions of genes (Extended Data Figure 7c) and found no evidence of off-target activity in *EPOR*-sg1-edited cells when compared to mock edited cells (Extended Data 7d-e).

### Integration of *tEPOR* cDNA at safe-harbor locus recapitulates the proliferative effect

Given the therapeutic utility of transgene integration at safe harbor sites^28^, we hypothesized that integration of a *tEPOR* cDNA at a safe harbor site may also enable increased erythroid production from edited HSPCs while leaving the endogenous *EPOR* locus intact. Given the fact that integration at the *CCR5* locus is an established method for delivery of therapeutic transgenes in HSPCs^29^, we developed a custom AAV6-packaged DNA repair template that would facilitate integration of an exogenous human UbC promoter driving expression of *tEPOR* cDNA followed by a T2A-YFP-BGH reporter (Figure 2a). Given the strong, constitutive expression of the UbC promoter, this method of insertion is expected to express *tEPOR* ubiquitously in all hematopoietic cell types, regardless of lineage.

**Figure 2:**
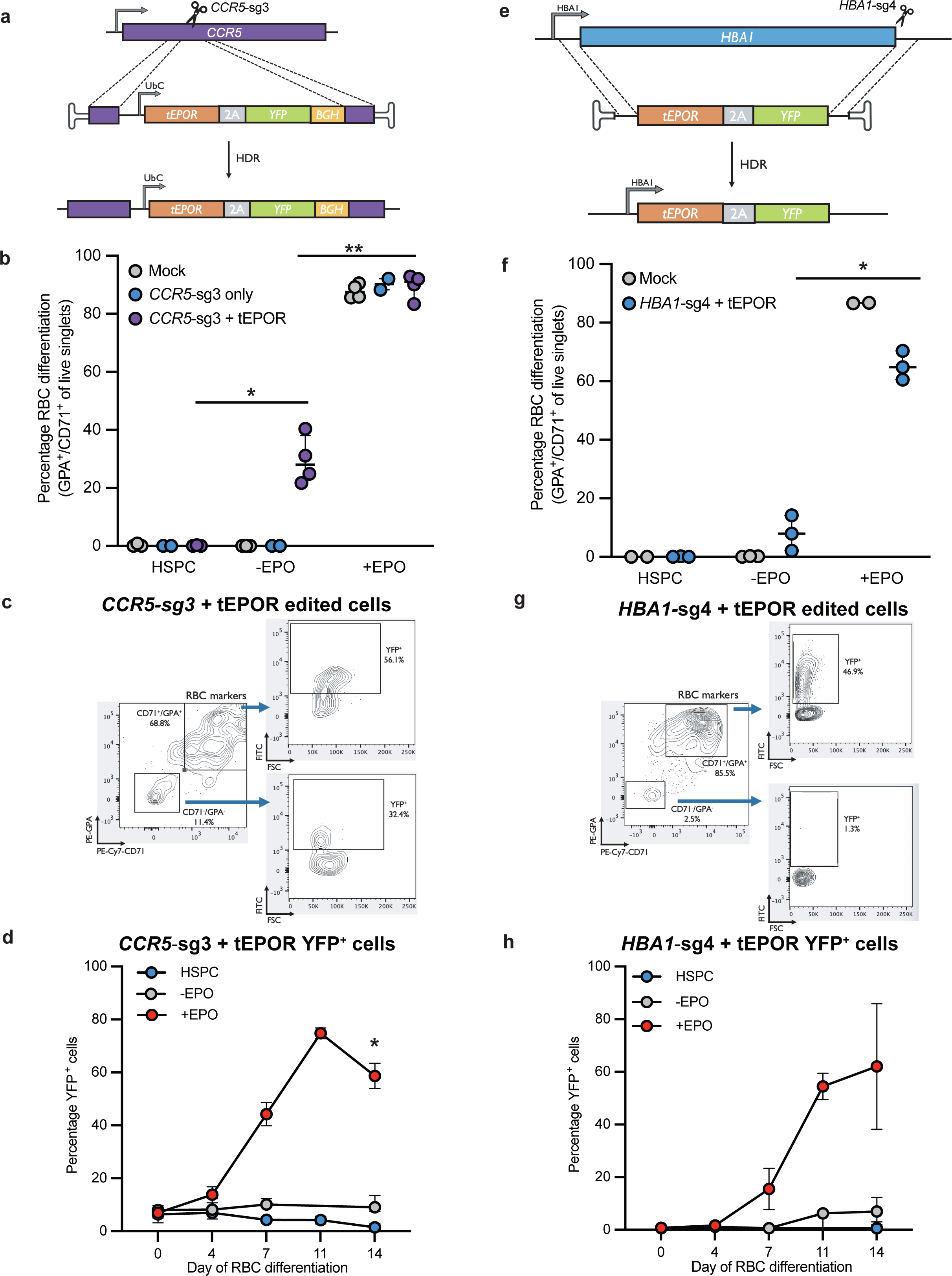
Integration of *tEPOR* cDNA demonstrates an erythroid-specific proliferative effect. **a.** Genome editing strategy to introduce *tEPOR*-T2A-YFP-BGH-polyA cDNA at the *CCR5* locus with expression driven by a ubiquitous UbC promoter. **b.** Percentage of GPA^+^/CD71^+^ of live single cells on day 14 of differentiation following introduction of *tEPOR* at *CCR5* locus. Bars represent median ± interquartile range. Values represent biologically independent HSPC donors: N=2-3 for HSPC condition, N=2-4 for -EPO and +EPO conditions. *P=0.0018 of -EPO vs. HSPC conditions; **P<0.0001 of -EPO vs. +EPO conditions by unpaired two-tailed *t*-test. **c.** Representative flow cytometry plots of one donor of *CCR5*-sg3 + tEPOR-edited HSPCs on day 14 of RBC differentiation in the +EPO condition. **d.** Percentage of YFP^+^ cells of live single cells as determined by flow cytometry. Points represent mean ± SEM. Values represent biologically independent HSPC donors: N=2 for HSPC condition, N=3 for -EPO condition, N=3-4 for +EPO condition. *P = 0.0003 of day 0 vs. day 14 by unpaired two-tailed *t*-test. **e.** Genome editing strategy to introduce *tEPOR*-T2A-YFP cDNA at the *HBA1* locus by whole gene replacement to place integration cassette under regulation of the endogenous *HBA1* promoter. **f.** Percentage of GPA^+^/CD71^+^ of live single cells on day 14 of differentiation following introduction of *tEPOR* cDNA at the *HBA1* locus. Bars represent median ± 95% confidence interval. Values represent biologically independent HSPC donors: N=2 for HSPC condition, N=2-3 for -EPO and +EPO condition. **\***P=0.0002 of -EPO to +EPO condition by unpaired two-tailed *t*-test. **g.** Representative flow cytometry plots of one donor of *HBA1*-sg4 + tEPOR-edited HSPCs on day 11 of RBC differentiation in the +EPO condition. **h.** Percentage of YFP^+^ cells of live single cells as determined by flow cytometry. Points represent mean ± SEM. Values represent biologically independent HSPC donors: N=2 for HSPC condition, N=3 for -EPO and +EPO condition.

To test this strategy for *tEPOR* expression, we edited HSPCs with Cas9 complexed with an established sgRNA targeting exon 2 of *CCR5*^29^ (*CCR5*-sg3), immediately followed by transduction with our custom DNA repair template. We then performed RBC differentiation post-editing and analyzed the kinetics of editing frequency, YFP expression, and erythroid differentiation using droplet digital PCR (ddPCR) and flow cytometry. As with endogenous *EPOR* truncation strategies, we observed more efficient erythroid differentiation in all treatments with EPO compared to the minus EPO conditions (Figure 2b). While flow cytometry confirmed the ubiquitous expression of YFP in edited cells, regardless of presence of CD71 and GPA erythroid markers during differentiation (Figure 2c), we did observe significant enrichment of YFP-expressing RBCs in the presence of EPO (P<0.0001 when comparing day 0 to day 14 of RBC differentiation (Figure 2d)). This enrichment was confirmed at the genomic level by ddPCR which showed an increase in percent edited alleles when *tEPOR*-expressing cells were subjected to erythroid differentiation, increasing by an average of 7.3-fold in the presence of EPO, as well as enrichment to a limited degree enrichment in the -EPO condition (Extended Data Figure 8a). With this editing strategy, we observed increased RBC production in the CCR5-sg3 + *tEPOR* condition and no increase in proliferation of CCR5-sg3 alone compared to mock control cells in the +EPO condition. We saw no increased proliferation in the cells maintained in HSPC media or in the -EPO condition (Extended Data Figure 8b). We then assessed if enrichment of edited cells differed along a gradient of EPO levels during RBC differentiation and again found that there were minimal differences in YFP^+^ cells in the 1, 3, and 20 U/mL conditions and a minor decrease in the 0.3 U/mL EPO condition compared to 0 U/mL EPO (Extended Data Figure 8c-d). There were comparable levels of RBC differentiation in all conditions except 0 U/mL EPO (Extended Data Figure 8e).

We once again observed that edited and unedited cells had no noticeable difference in lineage commitment or colony-forming ability following a colony-forming unit assay (Extended Data Figure 9a). In addition, we analyzed RBCs post-differentiation using hemoglobin tetramer HPLC and found that UbC-mediated expression of *tEPOR* resulted in both HgbF and HgbA expression but a relative increase in HgbF expression (Extended Data Figure 9b). These results indicate that expression of *tEPOR* from a safe harbor site is an effective means of driving increased RBC production from genome-edited HSPCs.

### Integration of *tEPOR* cDNA at *HBA1* demonstrates an erythroid-specific proliferative effect

While integration at a safe harbor locus effectively increased erythropoietic output from edited HSPCs, there is the concern that constitutive expression in all cell types could disrupt stem-ness or lead to other unintended effects. As an alternative, we can introduce promoter-less transgenes into endogenous genes in order for the integration cassette to be regulated by endogenous expression machinery. For instance, in prior work we designed a genome editing strategy to fully replace the *HBA1* gene with an *HBB* transgene in order to correct β-thalassemia^14^. We found that because α-globin is produced by duplicate genes, *HBA1* may serve as a safe harbor site to deliver custom payloads with strong erythroid-specific expression.

We therefore hypothesized that integration of the *tEPOR* cDNA at the *HBA1* locus could further enhance production of edited RBCs while avoiding potential complications with ubiquitous transgene expression because of the specificity of expression of *HBA1* in the RBC lineage. To test this hypothesis, we designed a custom integration cassette (also packaged in AAV6) to use with an established sgRNA (*HBA1*-sg4) to introduce a promoter-less *tEPOR* cDNA followed by a T2A-YFP reporter under expression of endogenous *HBA1* regulatory machinery (Figure 2e). Following editing, we observed efficient erythroid differentiation in all treatments with EPO and little to no differentiation in the absence of EPO (Figure 2f). We found that this integration strategy indeed yielded RBC-specific expression of YFP, which was only detectable as cells gained CD71 and GPA erythroid markers (Figure 2g). Over the course of RBC differentiation, we observed dramatic enrichment of YFP^+^ cells exclusively in the +EPO condition (Figure 2h). While we observed a mild degree of enrichment of edited alleles using ddPCR in the absence of EPO, this effect was more pronounced in the presence of EPO, eliciting an average 4.4-fold increase in percentage of edited alleles over the course of RBC differentiation (Extended Data Figure 10a). We again cultured edited cells in a gradient of EPO during RBC differentiation and this time observed minimal differences in YFP^+^ cell enrichment in all conditions containing EPO (Extended Data Figure 10b-c). There were also comparable levels of RBC differentiation in all conditions except 0 U/mL EPO (Extended Data Figure 10d).

Once again, we found that cells edited with *tEPOR* at *HBA1* demonstrated similar colony-forming ability as mock edited cells (Extended Data Figure 11a). Additionally, we found that *tEPOR*-expressing cells produce ratios of human hemoglobin that are similar to unedited RBCs (Extended Data Figure 11b).

### Combined *HBB*-*tEPOR* bicistronic cassettes simultaneously correct β-thalassemia and increase production of edited RBCs

Because the above results demonstrate that *tEPOR* expression yields increased RBC production from edited HSPCs, we then sought to couple this selective advantage with a therapeutic gene edit. We chose to combine tEPOR with our prior β-thalassemia correction approach to simultaneously correct the disease and increase production of clinically meaningful RBCs from these corrected HSPCs. One way this can be accomplished is by creating a bicistronic cassette that links expression of the therapeutic full-length *HBB* transgene with a *tEPOR* cDNA. Prior studies have found that the type of linker domain used can have a great bearing on transgene expression and protein function—particularly when function is dependent on formation of protein complexes, as is the case with the globin genes^14^. Therefore, we designed and tested a variety of AAV6 repair template vectors linking the two genes using standard T2A peptides, optimized T2A peptides with Furin cleavage sites^30^ (referred to as “FuT2A”), internal ribosome entry sites (referred to as “IRES”), as well by driving *tEPOR* expression from a separate exogenous promoter—human PGK1 (referred to as “PGK”). To evaluate the different vectors, we edited healthy donor HSPCs as previously described at *HBA1* using the bicistronic AAV6 repair templates and evaluated their ability to differentiate and enrich for edited RBCs over the course of erythroid differentiation (Figure 3a). While efficient RBC differentiation was achieved in all editing conditions (Figure 3b), we found that all four bicistronic cassettes drove >2-fold enrichment of edited alleles (range: 2.1-fold to 3.5-fold) (P=0.003 for PGK-tEPOR, P=0.0055 for T2A-tEPOR, P=0.0259 for F2A-tEPOR, and P=0.0003 for IRES-tEPOR when comparing day 0 to day 14) (Figure 3c-d). We note that allele targeting frequencies of 60% translates into >80% of the cells having at least one allele targeted (cell targeting frequency) and thus would not expect to see much more enrichment than we observed.

**Figure 3:**
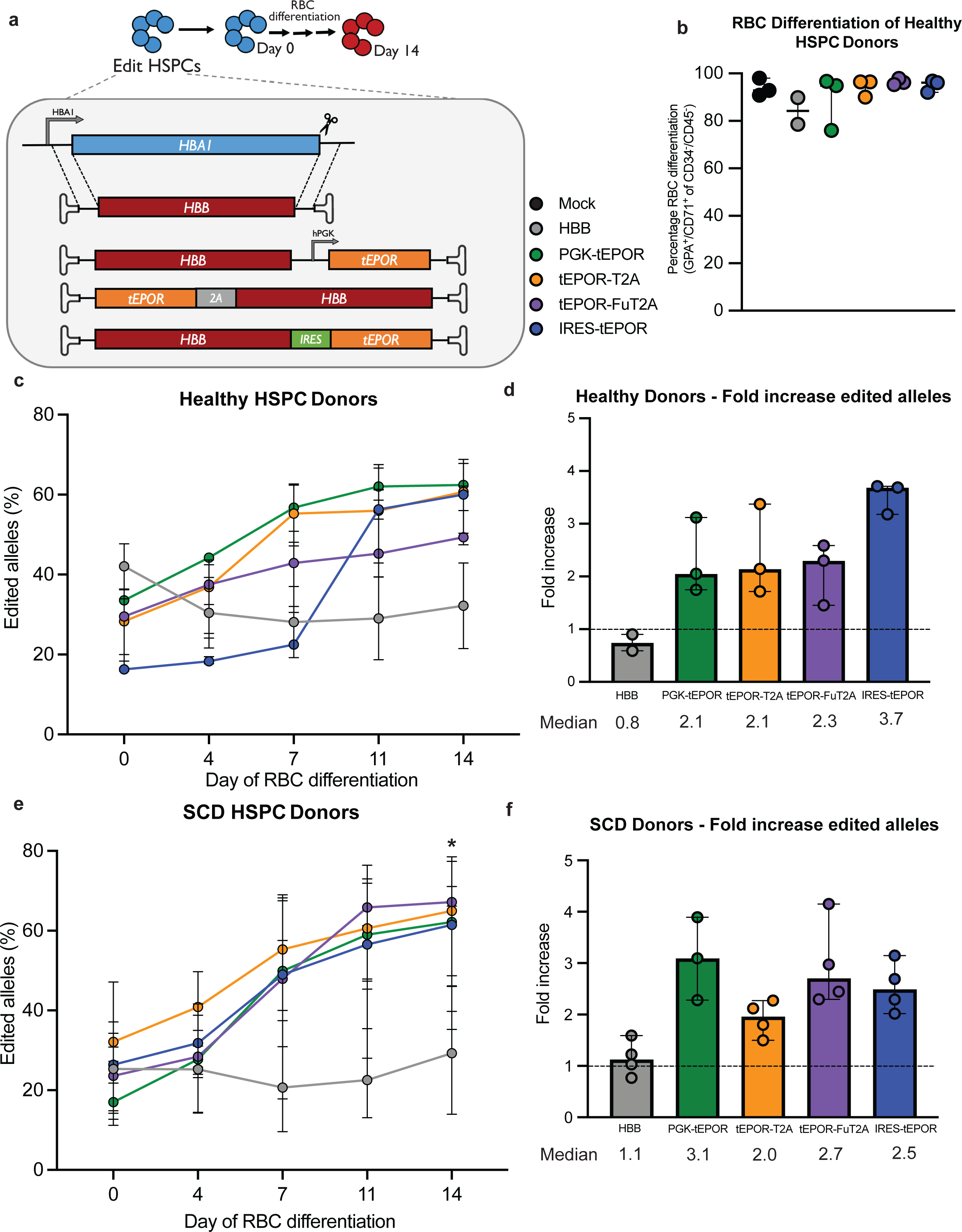
Therapeutic editing frequencies are achieved using bicistronic *HBB-tEPOR* cassette. **a.** Design of *HBB* (control) and *HBB*-*tEPOR* (bicistronic) AAV6 donor cassettes targeted to the *HBA1* locus by whole gene replacement. **b.** Percentage of GPA^+^/CD71^+^ of CD34^-^/CD45^-^ cells on day 14 as determined by flow cytometry. Points shown as median ± 95% confidence interval. Values represent biologically independent HSPC donors: N=2 for HBB, N=3 for all other vectors. **c.** Percentage of edited alleles for control (*HBB*) and bicistronic *HBB*-*tEPOR* in cord-blood derived CD34^+^ cells over the course of RBC differentiation. Points shown as median ± 95% confidence interval. Values represent biologically independent HSPC donors: N=2 for HBB, N=3 for all other vectors. **\***P=0.003 for PGK-*tEPOR*, P=0.0055 for T2A-*tEPOR*, P=0.0259 for Fu2A-*tEPOR*, and P=0.0003 for IRES-*tEPOR* (day 0 vs. day 14) by unpaired two-tailed *t*-test. **d.** Fold change in edited alleles from beginning (day 0) to end (day 14) of RBC differentiation. Bars represent median ± 95% confidence interval. **e.** Percentage of edited alleles for control (*HBB*) and bicistronic *HBB*-*tEPOR* vectors in sickle cell disease patient cells over the course of RBC differentiation. Points shown as median ± 95% confidence interval. Values represent biologically independent HSPC donors: N=3 for PGK-*tEPOR*, N=4 for all other vectors. **\***P=0.0061 for PGK-*tEPOR*, P=0.011 for T2A-*tEPOR*, P=0.0016 for Fu2A-*tEPOR*, P=0.0153 for IRES-*tEPOR* (day 0 vs. day 14) by unpaired two-tailed *t*-test. **f.** Fold change in edited alleles from beginning (day 0) to end (day 14) of RBC differentiation. Bars represent median ± standard deviation.

To ensure this strategy was also effective in patient-derived cells, we tested these bicistronic vectors in HSPCs derived from sickle cell disease (SCD) patients, this time comparing them to a therapeutic full-length *HBB* transgene^14^. All of the constructs are knocked-in to the *HBA1* locus without disrupting the endogenous *HBB* locus expressing HgbS. We found that vectors did not disrupt erythroid differentiation when compared to mock edited cells in the same donor, although their ability to differentiate was likely impacted by the variable quality of the frozen patient samples (Extended Data Figure 12a). Again, we observed >2-fold enrichment of edited alleles for all four bicistronic vectors (range: 2.0-fold to 3.1-fold), but no change in editing frequency for the original β-thalassemia correction vector (P=0.0061 for PGK-tEPOR, P=0.011 for T2A-tEPOR, P=0.0016 for Fu2A-tEPOR, P=0.0153 for IRES-tEPOR when comparing day 0 to day 14) (Figure 3e-f). We estimate that at the end of the differentiation almost all of the cells have at least one allele with the HBB-tEPOR knock-in and thus no biologic drive for further enrichment. When we analyzed differentiated RBCs for hemoglobin tetramers by HPLC, we found that the 2A vectors showed almost no HgbA expression (consistent with what has been seen previously in which addition of a 2A can disrupt HBB protein function^31^). In contrast, we found that PGK-tEPOR and IRES-tEPOR vectors showed an improvement in HgbA production relative to the *HBB*-only edited cells (Extended Data Figure 12b). There was no change in the HgbF expression in these samples.

### Multiplexed genome editing to introduce *tEPOR* and *HBB* leads to robust increase in *β-globin* mRNA in edited RBCs

In lieu of coupling the *tEPOR* cDNA and therapeutic edit at the same locus, an alternative strategy would be to multiplex two editing events at different loci to simultaneously truncate the endogenous *EPOR* and introduce the original β-thalassemia correction vector at *HBA1*. This may have the additional advantage that the endogenous *EPOR* truncation will more reliably recapitulate congenital erythrocytosis. We hypothesized that we could simultaneously deliver Cas9 separately pre-complexed with *EPOR*-truncating *EPOR*-sg1 and *HBA1*-sg4 sgRNA and then transduce HSPCs with both the β-thalassemia correction vector as well as the W439X *EPOR*-targeting vector. Because homology arms of each vector are specific for each site—*HBB* for the *HBA1* locus and W439X for the *EPOR* locus—integration for each vector will occur only at the intended locus. This strategy may allow simultaneous correction of β-thalassemia and increased erythropoietic output from corrected HSPCs. For this to be maximally effective, the two editing events must be present in the same cell. Therefore, during editing we used a DNA-PKcs inhibitor to increase the frequency of template integration at each locus^32,33^. Following editing, we determined whether this multiplexed editing strategy increases the frequency of corrected RBCs over the course of erythroid differentiation.

To model the clinical setting where both edited and unedited HSPCs would occupy the patient’s bone marrow, we introduced unedited cells at various concentrations at the start of erythroid differentiation (Figure 4a). Importantly, none of the multiplexed conditions disrupted erythroid differentiation compared to single-edited *HBB* or mock conditions (Figure 4b). In all of the multiplexed conditions, we observed an increase in the frequency of corrected alleles over the course of RBC differentiation (P=0.0332 for *HBB+tEPOR* 100%, P=0.0086 for *HBB+tEPOR* 30%, P=0.0122 for *HBB+tEPOR* 10% when comparing day 0 to day 14) (Figure 4c). In fact, in both the 30% and 10% multiplexed conditions, we achieved a higher frequency of edited alleles by the conclusion of erythroid differentiation compared to single-edited conditions with unedited cells introduced at an equivalent concentration (P=0.0113 for *HBB* versus *HBB+tEPOR* 30% at day 14 and P=0.008 for *HBB* versus *HBB+tEPOR* 10% at day 14) (Figure 4d). We confirmed truncation at the *EPOR* locus measured by GFP^+^ cells at day 14 of RBC differentiation (Extended Data Figure 13a). We also observed a corresponding increase in *HBB* mRNA expression from the *HBA1* locus in the 30% and 10% multiplexed edited conditions on day 14 of RBC differentiation compared to single-edited conditions (Figure 4e), indicating an improved ability for multiplexed editing to increase therapeutic potential of this β-thalassemia correction strategy. When we measured the colony-forming ability of the edited cells and found that differentiation into the various lineages was similar between mock cells and cells edited with *HBB* alone or *HBB+tEPOR*, although we did observe a decrease in total colonies produced in both edited conditions as would be expected from the increased amount of AAV used to target two loci^34,35^ (Extended Data Figure 13b-c).

**Figure 4:**
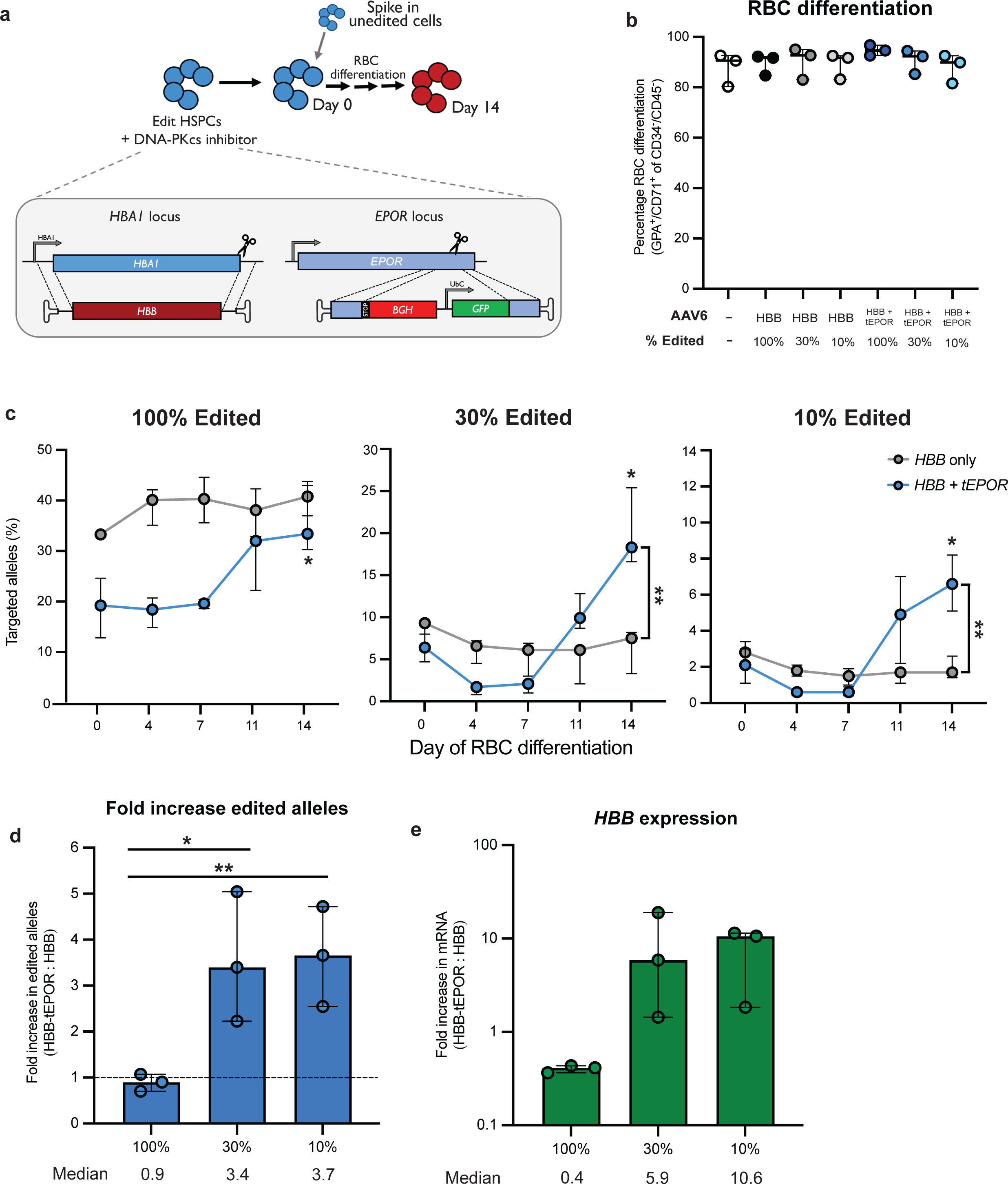
Multiplexed editing of *EPOR* and *HBA1* leads to robust increase in *HBB* mRNA within editing HSPCs. **a.** Schematic of multiplexed editing strategy with spike-in of unedited cells at start of erythroid differentiation to model HSPC transplantation. **b.** Percentage of GPA^+^/CD71^+^ of CD34^-^/CD45^-^ cells on day 14 of RBC differentiation as determined by flow cytometry. Points shown as median ± interquartile range. N=3 biologically independent HSPC donors. **c.** Percentage of edited alleles at *HBA1* in all multiplexed editing/spike-in conditions throughout RBC differentiation. Points shown as median ± 95% confidence interval. N=3 biologically independent HSPC donors. *P=0.0332 for *HBB*+*tEPOR* 100%, P=0.0086 for *HBB*+*tEPOR* 30%, P=0.0122 for *HBB+tEPOR* 10% (day 0 vs. day 14) by unpaired two-tailed *t*-test. **P=0.0113 for *HBB* versus *HBB*+*tEPOR* 30% at day 14 and P=0.008 for HBB versus *HBB*+*tEPOR* 10% at day 14 by unpaired two-tailed *t*-test. **d.** Fold increase in edited alleles on day 14 of differentiation of multiplexed conditions versus *HBB* only. Bars represent median ± 95% confidence interval. *P=0.0315 and **P=0.0123 by unpaired two-tailed *t*-test. **e.** mRNA expression of integrated *HBB* at *HBA1* locus normalized to *HBB* expression from mock. *GPA* mRNA expression was used as a reference. N=3 biologically independent HSPC donors. Bars represent median ± 95% confidence interval.

## Discussion

While insights from clinical genetics have typically implicated novel genes and pathways in disease, here we sought to use human genetics to develop novel strategies to *treat* disease. We used the precision of genome editing to capitalize on a previously characterized disorder called congenital erythrocytosis, which leads to EPO hypersensitivity and hyperproduction of erythrocytes, without causing pathology^4^. As demonstrated previously with variants identified in human genetics, such as those found in *CCR5*, *PCSK9*, and the gamma-globin promoter region, we hypothesized that we could use CRISPR to introduce this natural *EPOR* variant (*tEPOR*) to increase erythropoietic output from edited HSPCs. Prior work has highlighted the challenge of achieving long-term correction of disease following delivery of gene therapy or genome editing correction strategies^2,36,37^. While many efforts are underway to improve editing and engraftment frequencies, we hypothesized that we could develop a strategy to increase production of the clinically relevant cell type—the red blood cell—from edited HSPCs. If successful, then lower editing and engraftment frequencies could yield sufficient production of RBCs to achieve therapeutic benefit, and thus be curative for patients. Uchida *et al*. have shown that introducing the *tEPOR* variant using lentiviral delivery enhanced the efficacy of shRNA knockdown of *BCL11A* in upregulating HgbF, confirming the beneficial function of this variant^9^. However, our work deploys the precision of genome editing to generate tEPOR which may have broad utility across a spectrum of blood disorders.

We explored multiple genome-editing strategies to create the *tEPOR* variant, either through truncation of the endogenous *EPOR* gene or integration of a *tEPOR* cDNA at safe harbor loci. We found that HSPCs expressing *tEPOR* consistently demonstrated increased erythropoiesis but otherwise normal, EPO-dependent production of hemoglobin. In order to increase RBC production of genome-edited HSPCs in the context of disease correction, we combined the *tEPOR* cassette with a previously described β-thalassemia strategy in which the *HBB* gene replaces the *HBA1* gene using homology-directed repair-based genome editing^14^. This allowed us to simultaneously introduce an *HBB* transgene to restore normal hemoglobin production and to increase erythropoietic output from edited HSPCs. To demonstrate the flexibility of the various *tEPOR*-introduction strategies, we also developed an alternative multiplexed, site-specific genome editing strategy to pair the original β-thalassemia correction strategy with introduction of the *EPOR* truncation at the endogenous locus. Both strategies led to enrichment of genome-edited RBCs over the course of erythroid differentiation compared to the traditional β-thalassemia correction strategy. Because we found the effects to be EPO-dependent, it is possible the selective advantage *in vivo* may be even more pronounced because patients suffering from the hemoglobinopathies display elevated EPO levels due to their severe anemia^38^.

In terms of safety, the strategy to use CRISPR to introduce natural variants has the benefit of having been already “tested” *in vivo* in humans. However, it must be noted that several of the genome editing strategies introduce *tEPOR* under non-native regulation which could alter the normal function of tEPOR. In considering this possibility, we note while every cell in patients with CE harbor an *EPOR* truncation, therapeutic deployment of the genome editing strategy will result in the introduction of *tEPOR* in only a subset of HSPCs resident in the BM. Therefore, any aberrant effects of non-native *tEPOR* expression (e.g., bias away from lymphoid or other cell types) are unlikely to lead to cytopenia given the large number of unedited HSPCs remaining in the bone marrow post-transplant. Furthermore, by increasing erythropoietic output from edited HSPCs, we believe this work could enable the reduction or elimination of high-morbidity myeloablation regimens that are currently required to attain therapeutic levels of edited HSPCs. Expression of *tEPOR* could therefore be integrated into any treatment for blood disorders that involve transplantation of HSPCs. For example, even in an allogeneic hematopoietic stem cell transplant for red blood cell disorders, a truncation in the natural *EPOR* could be created using indel-based genome editing to give the derived transplanted erythroid progenitors a selective advantage. This strategy could thereby enable less toxic myeloablative conditioning to be effectively utilized where mixed chimerism might be the result.

Taken together, these results demonstrate the power of combining knowledge from human genetics with the precision of CRISPR genome editing technology to introduce clinically meaningful variants. As human genome sequencing becomes more commonplace and clinically routine^39,40^, it is likely that a greater number of variants of unknown significance will be discovered and characterized. We therefore believe that the strategy defined in this work—using CRISPR to introduce natural human variants—may be deployed to amplify the therapeutic potential of current and future cell therapies.

## Methods

### AAV6 vector design, production, and purification

Adeno-associated virus, serotype 6 (AAV6) vector plasmids were cloned into the pAAV-MCS plasmid (Agilent Technologies, Santa Clara, CA, USA), comprised of inverted terminal repeats (ITRs) derived from AAV2. Gibson Assembly Mastermix (New England Biolabs, Ipswich, MA, USA) was used for the creation of all DNA repair vectors as per manufacturer’s instructions. AAV6 vector was produced and purified with little variation from previously described processes^41^. 293T cells (Life Technologies, Carlsbad, CA, USA) were seeded in five 15 cm^2^ dishes with 13-15×10^6^ cells per plate 24 hours pre-transfection. Then, each dish was transfected with a standard polyethylenimine (PEI) transfection of 6 μg ITR-containing plasmid and 22 μg pDGM6 (gift from David Russell, University of Washington, Seattle, WA, USA), which holds the AAV6 cap, AAV2 rep, and Ad5 helper genes. Following a 48-72h incubation, cells were harvested and vectors were purified using the AAVpro purification kit (cat.: 6666; Takara Bio, Kusatsu, Shiga, Japan) as per manufacturer’s instructions and stored at -80 °C until further use. AAV6 vectors were titered using ddPCR to measure the number of vector genomes as previously described^42^.

### *In vitro* culture of CD34^+^ HSPCs

Human CD34^+^ HSPCs were cultured in conditions as previously described^13,43–46^. CD34^+^ HSPCs were isolated from cord blood (provided by Stanford Binns Program for Cord Blood Research) or sourced from Plerixafor- and/or G-CSF-mobilized peripheral blood (AllCells, Alameda, CA, USA and STEMCELL Technologies, Vancouver, Canada). Frozen Plerixafor- and/or G-CSF-mobilized peripheral blood of patients with sickle cell disease were provided by Dr. Vivien Sheehan, Emory University. CD34^+^ HSPCs were cultured at 1×10^5^–5×10^5^ cells/mL in StemSpan Serum-Free Expansion Medium II (STEMCELL Technologies, Vancouver, Canada) or Good Manufacturing Practice Stem Cell Growth Medium (SCGM, CellGenix, Freiburg, Germany) supplemented with a human cytokine (PeproTech, Rocky Hill, NJ, USA) cocktail: stem cell factor (100 ng/mL), thrombopoietin (100 ng/mL), Fms-like tyrosine kinase 3 ligand (100 ng/ml), interleukin 6 (100 ng/mL), streptomycin (20 mg/mL), and penicillin (20 U/mL), and 35 nM of UM171 (cat.: A89505; APExBIO, Houston, TX, USA). The cell incubator conditions were 37°C, 5% CO_2_, and 5% O_2_.

### Electroporation-aided transduction of cells

The synthetic chemically modified sgRNAs used to edit CD34^+^ HSPCs were purchased from Synthego (Redwood City, CA, USA) or TriLink Biotechnologies (San Diego, CA, USA) and were purified by HPLC. These modifications are comprised of 2′-O-methyl-3′-phosphorothioate at the three terminal nucleotides of the 5′ and 3′ ends described previously^17^. The target sequence for the sgRNAs were as follows:

*EPOR* sgRNA (*EPOR*-sg1)

5′-AGCTCAGGGCACAGTGTCCA-3′

*EPOR* sgRNA (*EPOR*-sg2)

5′-GCTCCCAGCTCTTGCGTCCA-3′

*CCR5* sgRNA (*CCR5*-sg3)

5′-GCAGCATAGTGAGCCCAGAA-3′

*HBA1* sgRNA (*HBA1*-sg4)

5′-GGCAAGAAGCATGGCCACCG-3′

The HiFi Cas9 protein was purchased from Integrated DNA Technologies (IDT) (Coralville, Iowa, USA) or Aldevron (Fargo, ND, USA). Preceding electroporation, RNPs were complexed at a Cas9:sgRNA molar ratio of 1:2.5 at 25°C for 10-20 min. Next, CD34^+^ cells were resuspended in P3 buffer (Lonza, Basel, Switzerland) with complexed RNPs and subsequently electroporated using the Lonza 4D Nucleofector and 4D-Nucleofector X Unit (program DZ-100). Electroporated cells were then plated at 1×10^5^-5×10^5^ cells/mL in the previously described cytokine-supplemented media. Immediately following electroporation, AAV6 was dispensed onto cells at 2.5×10^3^-5×10^3^ vector genomes/cell based on titers determined by ddPCR. For multiplex editing experiments, in addition to the steps described above, cells were incubated with 0.5 μM of a DNA-PKcs inhibitor, AZD7648 (cat.: S8843; Selleck Chemicals, Houston, TX) for 24 hours, as previously described^32,33^.

### Allelic modification analysis using ddPCR

Edited HSPCs were harvested within 2-3 days post-electroporation and at each media change throughout erythrocyte differentiation and then analyzed for modification frequencies of the alleles of interest. To quantify editing frequencies, we created custom ddPCR primers and probes to quantify HDR alleles (using in-out PCR and probe corresponding to the expected integration event) compared to an established genomic DNA reference (REF) at the *CCRL2* locus^14^. QuickExtract DNA extraction solution (Biosearch Technologies, Hoddesdon, UK, cat.: QE09050) was used to collect genomic DNA (gDNA) input, which was then digested using BamHI-HF or HindIII-HF as per the manufacturer’s instructions (New England Biolabs, Ipswich, MA, USA). The percentage of targeted alleles within a cell population was measured with a Bio-Rad QX200 ddPCR machine and QuantaSoft software (v.1.7; Bio-Rad, Hercules, CA, USA) using the following reaction mixture: 1-4 μL gDNA input, 10 μL of ddPCR SuperMix for Probes (no dUTP) (Bio-Rad), primer/probes (1:3.6 ratio; IDT), and volume up to 20 μL with H_2_O. ddPCR droplets were then generated following the manufacturer’s instructions (Bio-Rad): 20 μL of ddPCR reaction, 70 μL of droplet generation oil, and 40 μL of droplet sample. Thermocycler (Bio-Rad) settings were as follows: 98°C (10 min), 94°C (30 s), 55.7-60°C (30 s), 72°C (2 min), return to step 2 × 40– 50 cycles, and 98°C (10 min). Analysis of droplet samples was then performed using the QX200 Droplet Digital PCR System (Bio-Rad). We next divided the copies/μL for HDR (%): HDR (FAM) / REF (HEX). The following primers and probes were used in the ddPCR reaction:

*CCR5 (for tEPOR-YFP construct)*

Forward Primer (FP): 5’-GGGAGGATTGGGAAGACA-3’

Reverse Primer (RP): 5’-AGGTGTTCAGGAGAAGGACA-3’

Probe: 5’-6-FAM/AGCAGGCATGCTGGGGATGCGGTGG/3IABkFQ-3’

*HBA1 (for tEPOR-YFP construct)*

FP: 5’-AGTCCAAGCTGAGCAAAGA-3’

RP: 5’-ATCACAAACGCAGGCAGAG-3’

Probe: 5’-CGAGAAGCGCGATCACATGGTCCTGC-3’

*HBA1 (for HBB construct and tEPOR-HBB constructs)*

FP: 5’-GTGGCTGGTGTGGCTAATG-3’

RP: 5’-CAGAAAGCCAGCCAGTTCTT-3’

Probe: 5’-6-FAM/CCTGGCCCACAAGTATCACT/3IABkFQ-3’

*HBA1 (for HBB-tEPOR constructs)*

FP: 5’-TCTGCTGCCAGCTTTGAGTA-3’

RP: 5’-GCTGGAGTGGGACTTCTCTG-3’

Probe: 5’-6-FAM/ACTATCCTGGACCCCAGCTC/3IABkFQ-3’

*CCRL2 (reference)*

FP: 5’-GCTGTATGAATCCAGGTCC-3’,

RP: 5’-CCTCCTGGCTGAGAAAAAG -3’

Probe: 5’-HEX/TGTTTCCTC/ZEN/CAGGATAAGGCAGCTGT/3IABkFQ -3’

### Indel analysis using TIDE software

Within 2-4 days post-electroporation, HSPCs were harvested with QuickExtract DNA extraction solution (Biosearch Technologies, cat.: QE09050) to collect gDNA. The following primer sequences were used to amplify the respective cut site at the *EPOR* locus:

FP: 5’-CAGCTGTGGCTGTACCAGAA-3’

RP: 5’-CAGCCTGGTGTCCTAAGAGC-3’

Sanger sequencing of respective samples was then used as input for indel frequency analysis relative to a mock, unedited sample using TIDE as previously described^25^.

### *In vitro* differentiation of CD34^+^ HSPCs into erythrocytes

Following editing, HSPCs derived from healthy or sickle cell disease patients were cultured 2-3 days as described above. Subsequently, a 14-day *in vitro* differentiation was performed in supplemented SFEMII medium as previously described^24,47^. SFEMII base medium was supplemented with 100 U/mL penicillin–streptomycin, 10 ng/mL SCF (PeproTech, Rocky Hill, NJ, USA), 1 ng/mL IL-3 (PeproTech, Rocky Hill, NJ, USA), 3 U/mL EPO (eBiosciences, San Diego, CA, USA), 200 μg/mL transferrin (Sigma-Aldrich, St. Louis, MO, USA), 3% human serum (heat-inactivated from Sigma-Aldrich, St. Louis, MO, USA or Thermo Fisher Scientific, Waltham, MA, USA), 2% human plasma (isolated from umbilical cord blood provided by Stanford Binns Cord Blood Program), 10 μg/mL insulin (Sigma-Aldrich, St. Louis, MO, USA), and 3 U/mL heparin (Sigma-Aldrich, St. Louis, MO, USA). Cells were cultured in the first phase of medium for seven days at 1×10^5^ cells/mL. In the second phase of medium, day 7-10, cells were maintained at 1×10^5^ cells/mL, and IL-3 was removed from the culture. In the third phase of medium, day 11-14, cells were cultured at 1×10^6^ cells/mL, with a transferrin increase to 1 mg/mL.

### Immunophenotyping of differentiated erythrocytes

Differentiated erythrocytes were analyzed by flow cytometry on day 14 for erythrocyte lineage-specific markers using a FACS Aria II (BD Biosciences, San Jose, CA, USA). Edited and unedited cells were analyzed using the following antibodies: hCD45-V450 (HI30; BD Biosciences, San Jose, CA, USA), CD34-APC (561; BioLegend, San Diego, CA, USA), CD71-PE-Cy7 (OKT9; Affymetrix, Santa Clara, CA, USA), and CD235a-PE (GPA) (GA-R2; BD Biosciences). In addition to cell-specific markers, cells were also stained with Ghost Dye Red 780 (Tonbo Biosciences, San Diego, CA, USA) to measure viability.

### Hemoglobin tetramer analysis

Frozen pellets of approximately 1×10^6^ cells *in vitro*-differentiated erythrocytes were thawed and lysed in 30 µL of RIPA buffer with 1x Halt™ Protease Inhibitor Cocktail (Thermo Fisher Scientific, Waltham, MA, USA) for 5 minutes on ice. The mixture was vigorously vortexed and cell debris were removed by centrifugation at 13,000 RPM for 10 minutes at 4°C. HPLC analysis of hemoglobins in their native form was performed on a cation-exchange PolyCAT A column (35 × 4.6 mm^2^, 3 µm, 1,500 Å) (PolyLC Inc., Columbia, MD, USA) using a Perkin-Elmer Flexar HPLC system (Perkin-Elmer, Waltham, MA, USA) at room temperature and detection at 415 nm. Mobile phase A consisted of 20 mM Bis-tris and 2 mM KCN at pH 6.94, adjusted with HCl. Mobile phase B consisted of 20 mM Bis-tris, 2 mM KCN, and 200 mM NaCl at pH 6.55. Hemolysate was diluted in buffer A prior to injection of 20 µL onto the column with 8% buffer B and eluted at a flow rate of 2 mL/min with a gradient made to 40% B in 6 min, increased to 100% B in 1.5 min, and return to 8% B in 1 min and equilibrated for 3.5 min. Quantification of the area under the curve of the peaks was performed with TotalChrom software (Perkin-Elmer, Waltham, MA, USA) and raw values were exported to GraphPad Prism 9 for plotting and further analysis.

### mRNA analysis

Following differentiation of HSPCs into erythrocytes, cells were harvested, and RNA was extracted using the RNeasy Plus Mini Kit (Qiagen, Hilden, Germany). Subsequently, complementary DNA was made from approximately 100 ng of RNA using the iScript Reverse Transcription Supermix for quantitative PCR with reverse transcription (Bio-Rad, Hercules, CA, USA). Expression levels of β-globin transgene and α-globin mRNA were quantified with a Bio-Rad QX200 ddPCR machine and QuantaSoft software (v.1.7; Bio-Rad, Hercules, CA, USA) using the following primers and 6-FAM/ ZEN/IBFQ-labeled hydrolysis probes, purchased as custom-designed PrimeTime qPCR Assays from IDT:

*HBB and HBB-tEPOR into HBA1:*

FP: 5’-GGTCCCCACAGACTCAGAGA-3’

RP: 5’-CAGCATCAGGAGTGGACAGA

Probe: 5’-6-FAM/AACCCACCATGGTGCATCTG/3IABkFQ -3’

To normalize for RNA input, levels of the RBC-specific reference gene GPA were determined in each sample using the following primers and HEX/ZEN/IBFQ-labeled hydrolysis probes, purchased as custom-designed PrimeTime qPCR Assays from IDT: *GPA (reference):*

FP: 5′-ATATGCAGCCACTCCTAGAGCTC-3′

RP: 5′-CTGGTTCAGAGAAATGATGGGCA-3′

Probe: 5’-HEX/AGGAAACCGGAGAAAGGGTA/3IABkFQ -3’

ddPCR reactions were created using the respective primers and probes, and droplets were generated as described above. Thermocycler (Bio-Rad; settings were as follows: 98°C (10 min), 94°C (30 s), 54°C (30 s), 72°C (30 s), return to step 2 × 50 cycles, and 98°C (10 min). Analysis of droplet samples was done using the QX200 Droplet Digital PCR System (Bio-Rad). To determine relative expression levels, the numbers of *HBB* transgene copies/mL were divided by the numbers of *GPA* copies/mL.

### Methylcellulose colony forming unit (CFU) assay

2-3 days post-electroporation HSPCs were plated in SmartDish 6-well plates (cat.: 27370; STEMCELL Technologies, Vancouver, Canada) containing MethoCult H4434 Classic or MethoCult H4434 Classic without EPO (cat.: 04444; cat: 04544.; STEMCELL Technologies, Vancouver, Canada). After 14 days, the wells were imaged using the STEMvision Hematopoietic Colony Counter (STEMCELL Technologies, Vancouver, Canada). Colonies were counted and scored to determine the number of BFU-E, CFU-E, CFU-GM, and CFU-GEMM colonies.

### Quantification of editing efficiency at evaluated off-target sites

Potential sgRNA off-target sites were predicted using the CRISPR Off-target Sites with Mismatches, Insertions, and Deletions (COSMID) online tool^48^. Sites were ranked according to score and duplicate predictions at the same location were removed. All sites with a score ≤5.5 were included in analysis and the 5 sites in exonic or UTR regions were further analyzed. PCR amplification of these sites was performed using genomic DNA from mock-edited and RNP-edited cells. The following primers were used with Illumina adapters (FP adapter: 5’-ACACTCTTTCCCTACACGACGCTCTTCCGATCT-3’, RP adapter: 5’-GACTGGAGTTCAGACGTGTGCTCTTCCGATCT-3’):

*EPOR-OT1*

FP: 5’-GAGCGGGCTACAGAGCTAGA-3’

RP: 5’-TGGCAGAAAGTAAGGGGATG-3’

*EPOR-OT2*

FP: 5’-ACTTGTGGAGCCACAGTTTG-3’ RP: 5’-AATGCCCTTGAGATGAATGC-3’

*EPOR-OT3*

FP: 5’-TCACACACCCGTAGCCATAA-3’

RP: 5’-AGAATGCTCTTTGCAGTAGCC-3’

*EPOR-OT4*

FP: 5’-CTCAAAACTTCACCCAGGCT-3’

RP: 5’-GGTCTGTCATTGAATGCCTT-3’

*EPOR-OT5*

FP: 5’-CAACCCTGATGGGTCTGC-3’

RP: 5’-CCACAGCTGGCTGACCTT-3’

Following amplification, PCR products were purified by gel electrophoresis and subsequent extraction using the GeneJet Gel Extraction Kit (Thermo Fisher Scientific, Waltham, MA, cat.: FERK0692). Purified samples were submitted for library preparation and sequencing by Amplicon-EZ NGS (Azenta Life Sciences, San Francisco, CA), ensuring a yield of over 100,000 reads per sample. Amplicons, flanked by Illumina partial adapter sequences, which encompassed the programmed double-strand breaks (DSBs) for CRISPR/Cas9, underwent sequencing using Illumina chemistry. FastQC (v0.11.8, http://www.bioinformatics.babraham.ac.uk/projects/fastqc/, default parameters) was employed to assess the quality of raw reads. Subsequently, paired-end reads were aligned to the specified off-target regions using CRISPResso2 (version 2.2.14, CRISPResso --fastq_r1 reads_r1.fastq.gz --fastq_r2 reads_r2.fastq.gz -- amplicon_seq)^49^.

### Statistical analysis

GraphPad Prism 9 software was used for all statistical analysis.

### Data availability

All data supporting the findings of this study are available within the paper and its Supplementary Information. High-throughput sequencing data generated for off-target analysis will be uploaded to the NCBI Sequence Read Archive submission.

## Acknowledgements

The authors thank the following funding sources that made this work possible: American Society of Hematology Junior Faculty Scholar Award to M.K.C.; Stanford Medical Scholars Research Program, the American Society of Hematology Minority Medical Student Award Program, and the Stanford Medical Scientist Training Program to S.E.L.; American Society of Hematology Minority Medical Student Award Program to J.C. We would also like to thank the Stanford Binns Program for Cord Blood Research for providing CD34^+^ HSPCs and the FACS Core Facility at the Stanford Institute of Stem Cell Biology and Regenerative Medicine for access to FACS machines.

