## Supplemental Figures for "Using human genetics to develop strategies to increase erythropoietic output from genome-edited hematopoietic stem and progenitor cells"

#### Extended Data Figure 1

a

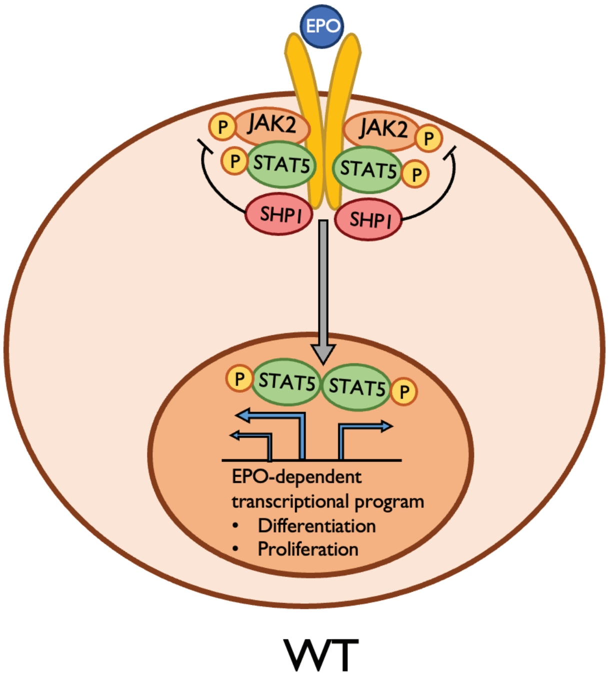

**Extended Data Figure 1. Schematic of wild-type (WT) and tEPOR signaling cascade.**

**a.** Visual representation of EPOR signaling cascade. SHP1 binds to intracellular inhibitory domain in WT cells, downregulating EPOR signaling. This domain is truncated in tEPOR cells making them hypersensitive to EPO.

### Extended Data Figure 2

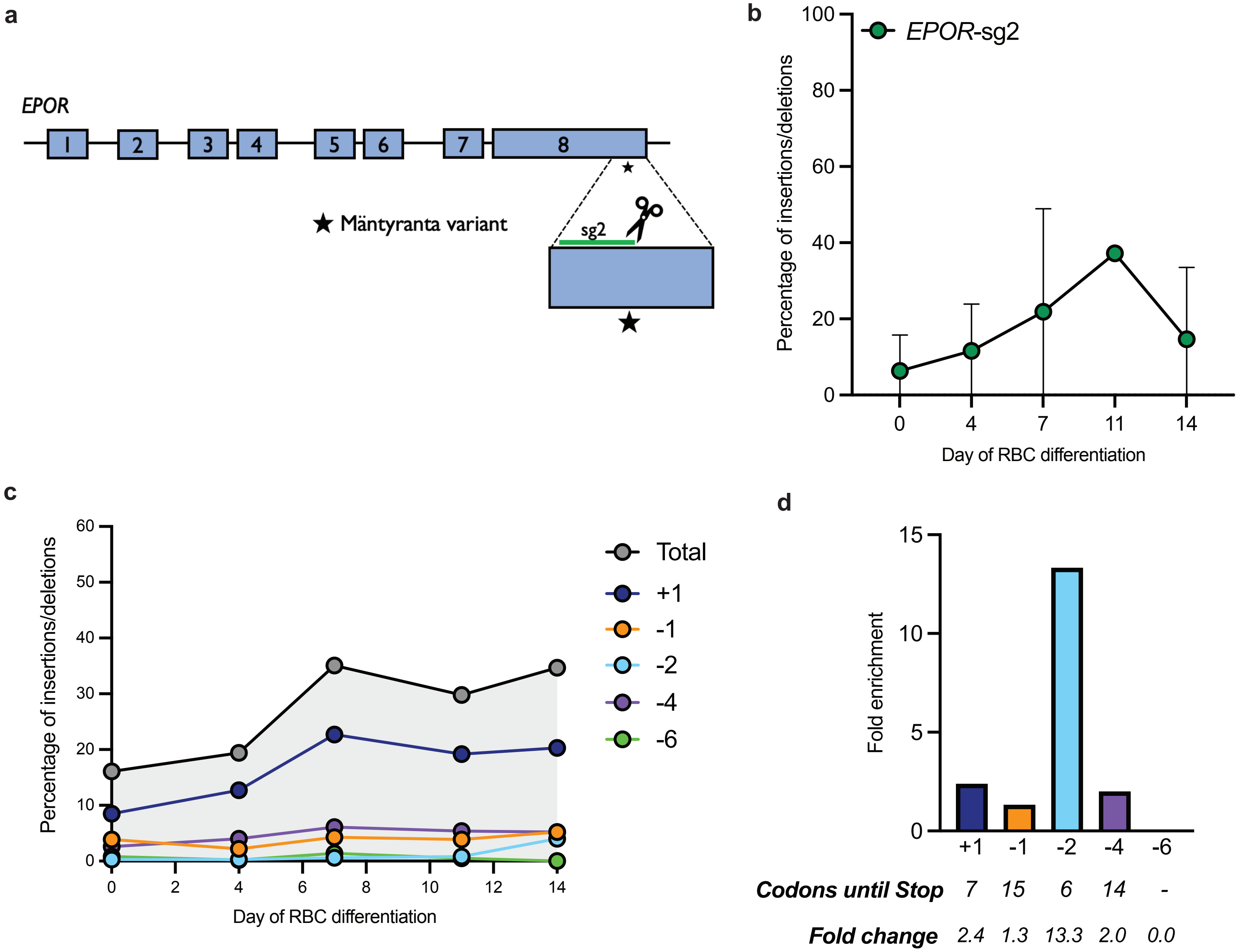

**Extended Data Figure 2. Indel analysis of HSPCs edited with second candidate *EPOR* sgRNA (sg2) over course of erythroid differentiation.**

**a.** Schematic of *EPOR* gene and location of the second candidate sgRNA (*EPOR*-sg2) indicated by a line. Location of c.1316G>A mutation is denoted by the star **b.** Frequency of indels created by *EPOR*-sg2 in primary human $CD34^+$ HSPCs over the course of erythroid differentiation. Points represent median ± interquartile range. Values represent N=4 biologically independent HSPC donors. **c.** Frequency of five most common indels found in one HSPC donor targeted with *EPOR*-sg2 over the course of RBC differentiation. **d.** Fold enrichment of five most common indels over course of RBC differentiation in one HSPC donor targeted with *EPOR*-sg2.

#### Extended Data Figure 3

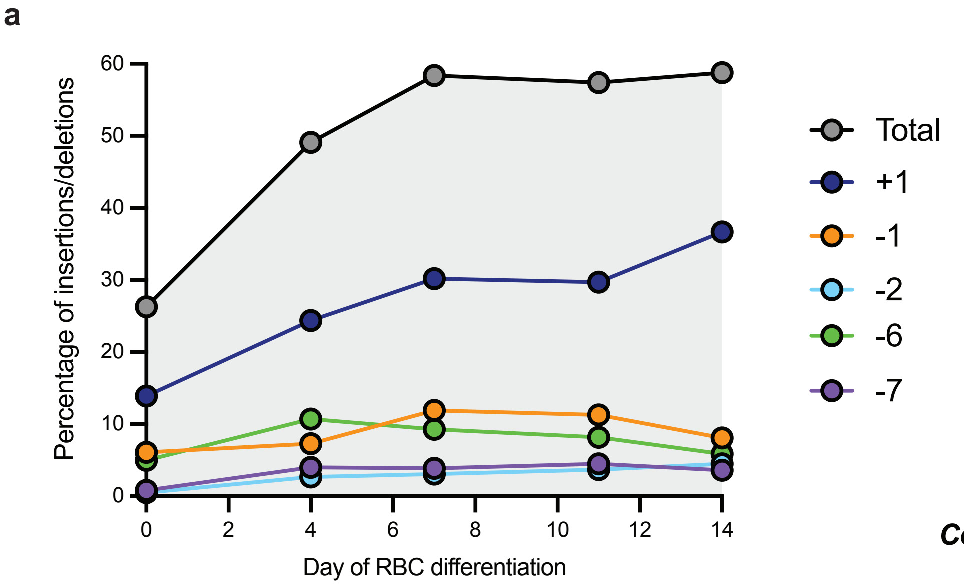


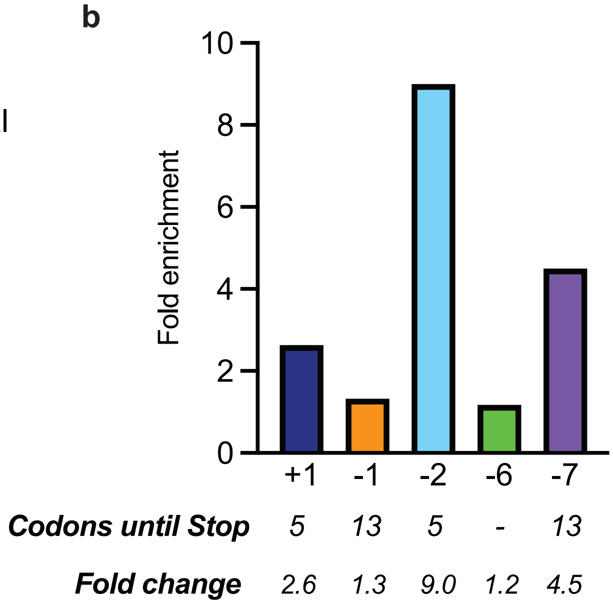

**Extended Data Figure 3. Indel analysis of HSPCs edited with most effective *EPOR* sgRNA (sg1) over course of erythroid differentiation.**

**a.** Frequency of five most common indels found in one HSPC donor targeted with *EPOR*-sg1over the course of RBC differentiation. **b.** Fold enrichment of five most common indels over course of RBC differentiation in one HSPC donor targeted with *EPOR*-sg1.

**Extended Data Figure 4**

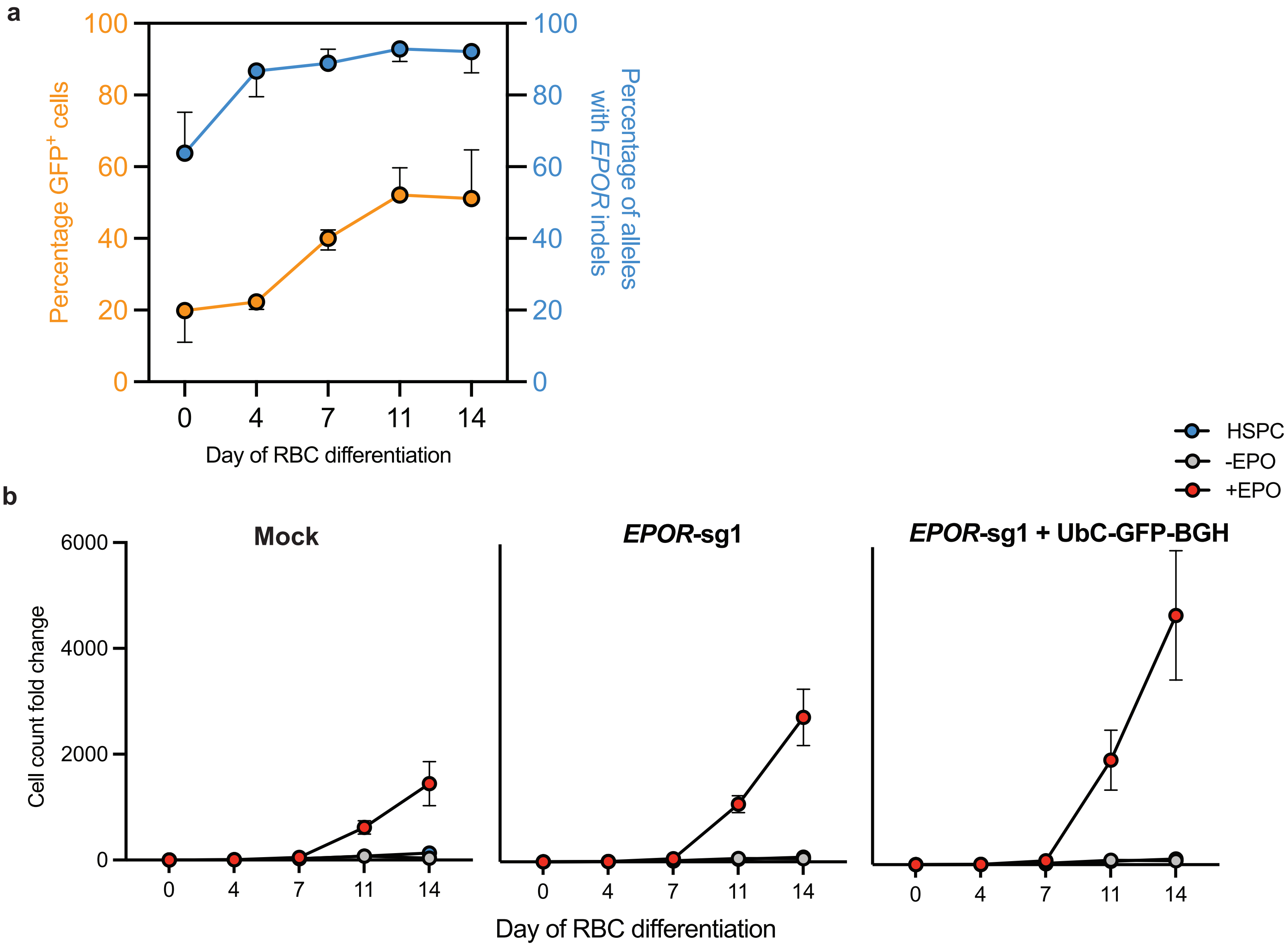

**Extended Data Figure 4. Almost all cells edited at *EPOR* locus contain tEPOR at end of differentiation and show increased erythroid proliferation.**

**a.** Plot of percentage of $GFP^+$ cells of live single cells maintained in RBC media +EPO (from same data shown in Figure 1e) overlaid with percentage of alleles containing indels in *EPOR* in same three biological donors. Points represent median ± 95% confidence interval. **b.** Cell count fold change in mock and edited cells maintained in RBC media +/- EPO or HSPC media. Points represent mean ± SEM. Values represent biologically independent HSPC donors N=3 for Mock and *EPOR*-sg1+ BGH and N=2 for sg1.

#### Extended Data Figure 5

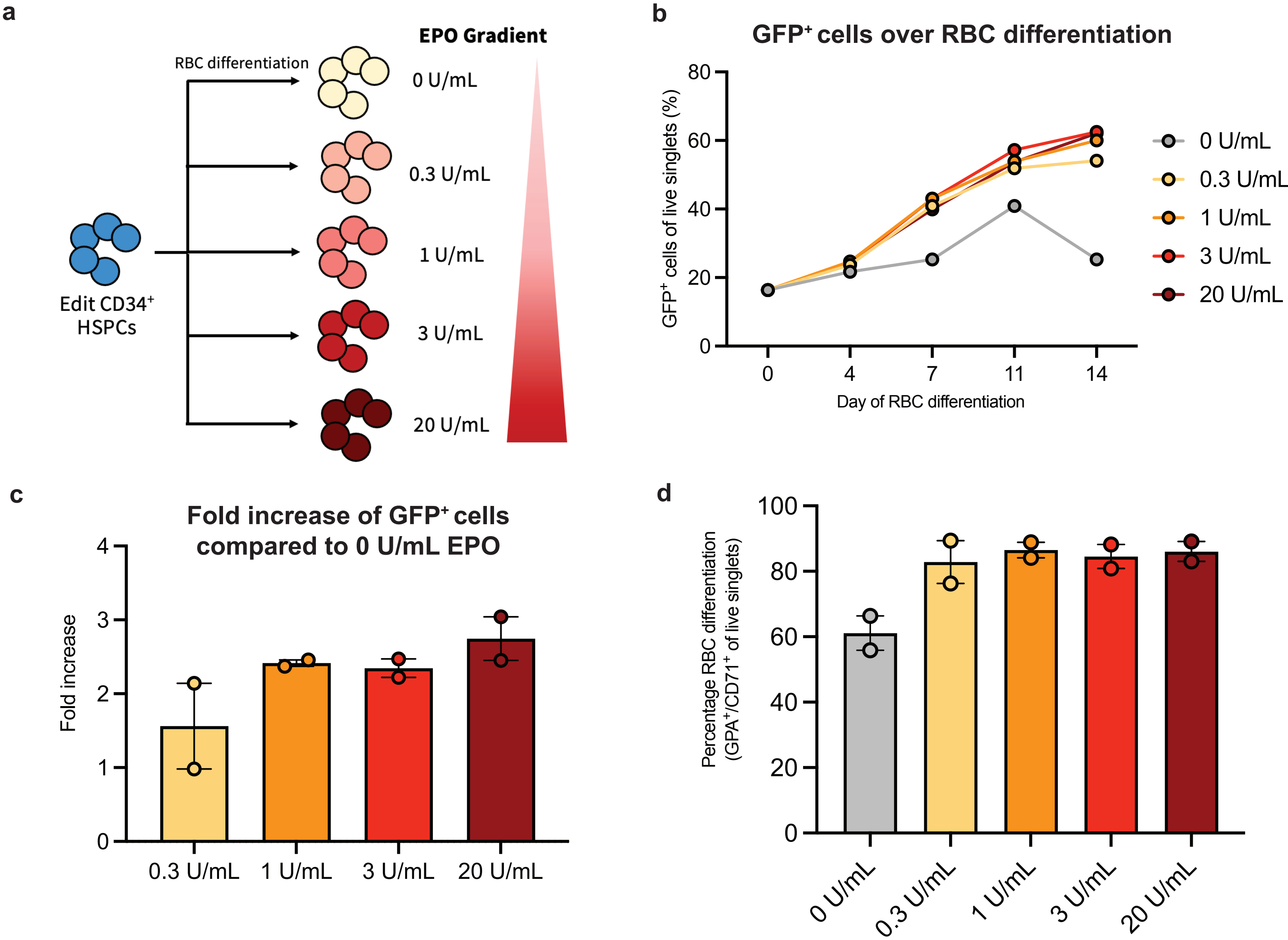

**Extended Data Figure 5. Enrichment of edited cells and RBC differentiation is only minorly affected by different concentration of EPO in media.**

**a.** Schematic depicting HSPC editing and subsequent RBC differentiation using different levels of EPO cytokine. **b.** Percentage of $GFP^+$ cells of live single cells maintained in RBC media with 0 U/mL – 20 U/mL EPO from one biological HSPC donor. **c.** Fold increase of $GFP^+$ cells at each concentration of EPO compared to 0 U/mL EPO at day 14 of RBC differentiation. Bars represent median ± 95% confidence interval. N=2 biological HSPC donors. **d.** Percentage of $GPA^+/CD71^+$ of live single cells on day 14 of differentiation. Bars represent median ± 95% confidence interval. Values represent N=2 biologically independent HSPC donors.

#### Extended Data Figure 6

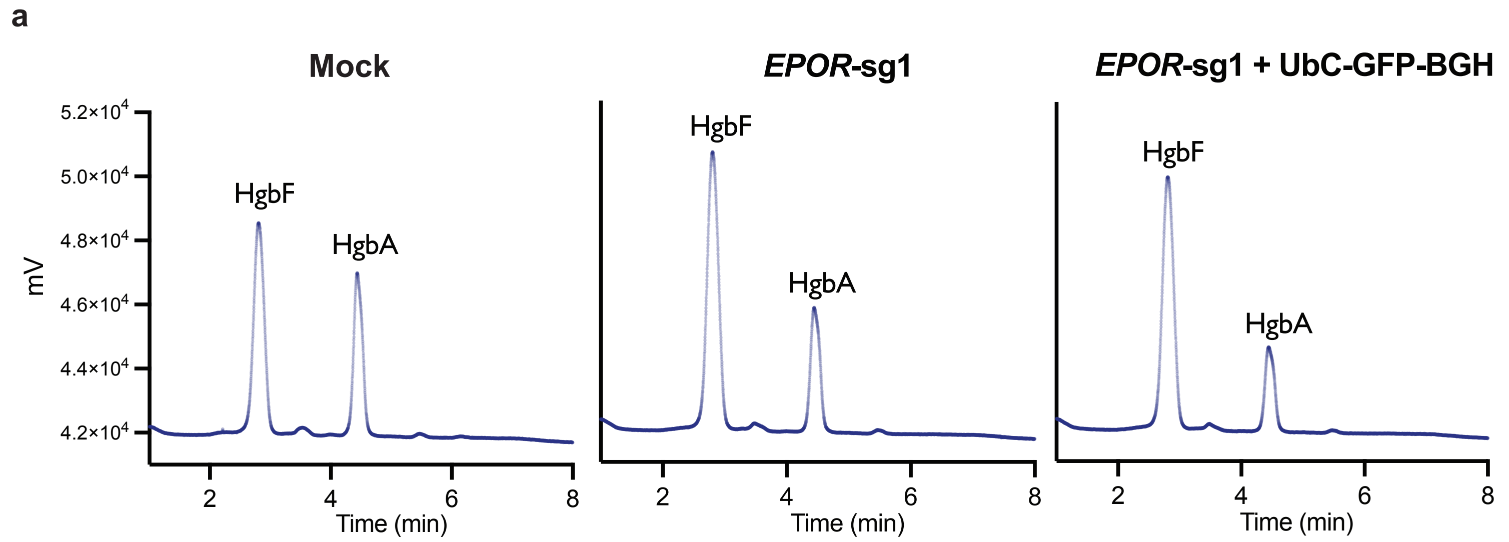


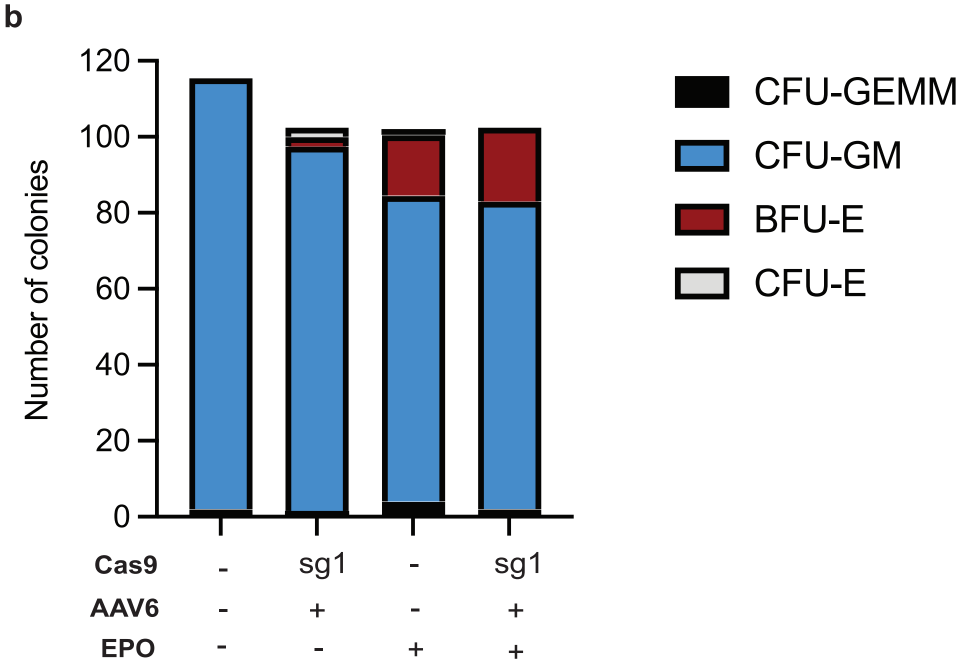

**Extended Data Figure 6. HSPCs edited at *EPOR* locus show normal hemoglobin tetramers and maintain lineage formation.**

**a.** Representative HPLC plots of cells targeted with *EPOR*-sg1 at day 14 of RBC differentiation. HgbF=fetal hemoglobin, HgbA=adult hemoglobin. **b.** Colony formatting unit (CFU) assay of mock edited cells versus cells edited with *EPOR*-sg1 + BGH*.* Bars represent total number of colonies of each type: CFU-GEMM (multi-potential granulocyte, erythroid, macrophage, megakaryocyte progenitor cells), CFU-GM (colony forming unit-granulocytes and monocytes), BFU-E (erythroid burst forming units), CFU-E (colony forming unit-erythroid) colonies. N=1.

### Extended Data Figure 7

**a**

#### Predicted sites with score ≤ 5.5

| | Chromosomal Position | Score | Region | Gene |
|---|---|---|---|---|
| On-target | Chr19:11378177-11378199 | 0 | Exon | *EPOR* |
| Off-target | Chr4:2834819-2834841 | 0.54 | 5' UTR | *SH3BP2* |
| | Chr17:83145508-83145530 | 0.61 | Intergenic | |
| | Chr14:74552967-74552989 | 0.63 | Exon | *LTBP2* |
| | Chr17:76190490-76190511 | 1.02 | Intron | *RNF157* |
| | Chr17:74795165-74795187 | 1.08 | Intron | *TMEM104* |
| | Chr19:57614583-57614605 | 1.22 | Intron | *ZNF134* |
| | Chr22:23681747-23681770 | 1.25 | Intron | *GUSBP11* |
| | Chr9:133742339-133742361 | 1.44 | Intergenic | |
| | Chr1:109777058-109777079 | 1.48 | Intergenic | |
| | Chr17:3725613-3725635 | 1.5 | Exon | *HASPIN* |
| | Chr20:47957087-47957110 | 1.51 | Intergenic | |
| | Chr15:62897167-62897188 | 1.52 | Intergenic | |
| | Chr1:153353786-153353807 | 1.53 | Intergenic | |
| | Chr22:47588374-47588397 | 1.79 | Intergenic | |
| | Chr5:66475949-66475972 | 1.82 | Intergenic | |
| | Chr15:35375033-35375054 | 1.83 | Intron | *DPH6* |
| | Chr3:146694160-146694183 | 2.08 | Intergenic | |
| | ChrX:53455142-53455163 | 2.15 | Intergenic | |
| | Chr16:58089238-58089259 | 2.15 | Intergenic | |
| | Chr15:66336597-66336618 | 2.15 | 3' UTR | *TIPIN* |
| | Chr11:69809008-69809031 | 2.17 | Intergenic | |
| | Chr8:125489832-125489853 | 2.21 | Intergenic | |
| | Chr8:1741386-1741407 | 2.35 | Intergenic | |
| | Chr6:74015621-74015642 | 2.35 | Intergenic | |
| | Chr2:207213231-207213252 | 2.66 | Intron | *SLID-AS1* |
| | Chr7:68423063-68423084 | 2.69 | Intergenic | |
| | Chr11:73258289-73258311 | 2.72 | Intergenic | |
| | Chr2:45355014-45355035 | 2.95 | Intergenic | |
| | Chr3:60743768-60743789 | 2.95 | Intron | *FHIT* |
| | Chr6:103967054-103967075 | 3.04 | Intergenic | |
| | Chr3:22886579-22886600 | 3.06 | Intergenic | |
| | Chr16:88736230-88736251 | 3.11 | Exon | *PIEZO1* |
| | Chr2:120715491-120715512 | 3.11 | Intergenic | |
| | ChrX:68993580-68993602 | 3.17 | Intergenic | |
| | Chr5:43348327-43348348 | 3.18 | Intergenic | |
| | Chr9:123359014-123359035 | 3.2 | Intron | *CRB2* |
| | ChrX:30077883-30077904 | 3.21 | Intergenic | |
| | Chr22:34432694-34432715 | 3.21 | Intergenic | |
| | Chr21:45947860-45947882 | 3.31 | Intergenic | |
| | Chr12:94243003-94243025 | 3.35 | Intron | *PLXNC1* |
| | Chr18:79252201-79252223 | 3.4 | Intron | *ATP9B* |
| | Chr8:144201447-144201469 | 3.42 | Intron | *MROH1* |
| | Chr2:3859090-3859111 | 3.48 | Intergenic | |
| | Chr5:23614195-23614216 | 3.56 | Intergenic | |
| | Chr10:44054693-44054715 | 3.62 | Intergenic | |
| | Chr3:59498371-59498393 | 3.62 | Intergenic | |
| | Chr22:48885862-48885883 | 3.63 | Intron | *LINC01310* |
| | Chr2:236161588-236161609 | 3.63 | Intergenic | |
| | Chr13:76821039-76821060 | 3.66 | Intergenic | |
| | ChrX:69384155-69384176 | 3.78 | Intergenic | |
| | Chr6:22043322-22043343 | 3.89 | Intron | *CACS15* |
| | Chr19:54221554-54221575 | 3.89 | Intron | *LILRB3* |
| | Chr19:54241302-54241323 | 3.89 | Intron | *LILRA6* |
| | Chr3:171095217-171095238 | 3.95 | Intron | *TNIK* |
| | Chr17:29069628-29069649 | 3.96 | Intergenic | |
| | Chr5:79849804-79849827 | 4.02 | Intergenic | |
| | Chr13:23653867-23653888 | 4.03 | Intron | *TNFRSF19* |
| | Chr6:31225407-31225428 | 4.03 | Intergenic | |
| | Chr17:3169786-3169807 | 4.09 | Intron | *LOC100288728* |
| | Chr1:209455360-209455383 | 4.12 | Intergenic | |
| | Chr22:42906036-42906057 | 4.2 | Intron | *PACSIN2* |
| | Chr10:48290891-48290913 | 4.36 | Intergenic | |
| | Chr9:136644277-136644299 | 4.41 | Intergenic | |
| | Chr19:57018808-57018830 | 4.5 | Intergenic | |
| | Chr22:25314961-25314982 | 4.66 | Intergenic | |
| | Chr20:39849896-39849917 | 4.8 | Intergenic | |
| | Chr9:1374116-1374137 | 4.82 | Intergenic | |
| | Chr1:223229923-223229944 | 4.88 | Intron | *SUSD4* |
| | Chr6:54602796-54602819 | 4.95 | Intergenic | |
| | Chr14:103074393-103074414 | 4.95 | Intergenic | |
| | Chr16:50151538-50151561 | 4.99 | Intergenic | |
| | Chr7:91541296-91541318 | 5 | Intergenic | |
| | Chr6:52211164-52211187 | 5.08 | Intergenic | |
| | Chr5:122639266-122639287 | 5.14 | intron | *LINC02201* |
| | Chr14:105666438-105666459 | 5.18 | Intergenic | |
| | Chr13:44678373-44678396 | 5.39 | intron | *LINC00407* |
| | Chr1:154810236-154810258 | 5.4 | intron | *KCNN3* |
| | Chr15:47975646-47975668 | 5.49 | Intergenic | |

**b**

#### OT distribution by region in genome

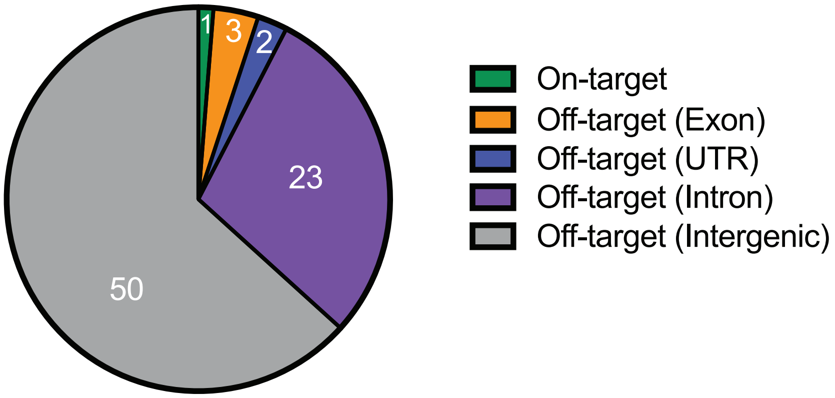


**c**

#### Exonic and UTR OT Sites

| Off-target Site | Chromosomal Position | COSMID Score | Sequence | Mismatch | Strand | Cut Site | Gene | Region of Gene |
|---|---|---|---|---|---|---|---|---|
| OT1 | Chr4:2834819-2834841 | 0.54 | TGCTTACGGCACAGTGTCCAGGG -- hit<br>AGCTCAGGGCACAGTGTCCANGG -- query | 3 | + | 2834835 | *SH3BP2* | 5' UTR |
| OT2 | Chr14:74552967-74552989 | 0.63 | AGCTGTTGGCACAGTGTCCACGG -- hit<br>AGCTCAGGGCACAGTGTCCANGG -- query | 3 | + | 74552983 | *LTBP2* | Exon |
| OT3 | Chr17:3725613-3725635 | 1.5 | AACTCAGTGCACTGTGTCCAGGG -- hit<br>AGCTCAGGGCACAGTGTCCANGG -- query | 3 | + | 3725629 | *HASPIN* | Exon |
| OT4 | Chr15:66336597-66336618 | 2.15 | A^CTCTGGGCACACTGTCCATGG -- hit<br>AGCTCAGGGCACAGTGTCCANGG -- query | 2 | + | 66336612 | *TIPIN* | 3' UTR |
| OT5 | Chr16:88736230-88736251 | 3.11 | AGCTCAGGGCGCAG^GTCCATGG -- hit<br>AGCTCAGGGCACAGTGTCCANGG -- query | 1 | + | 88736245 | *PIEZO1* | Exon |

**d**

#### NGS Read Depth

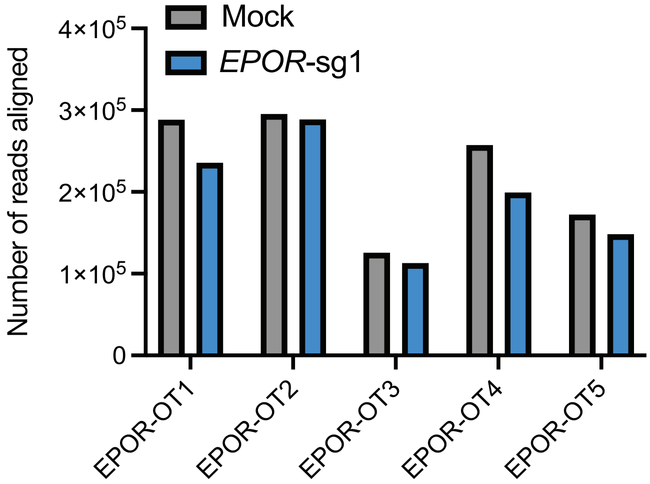


**e**

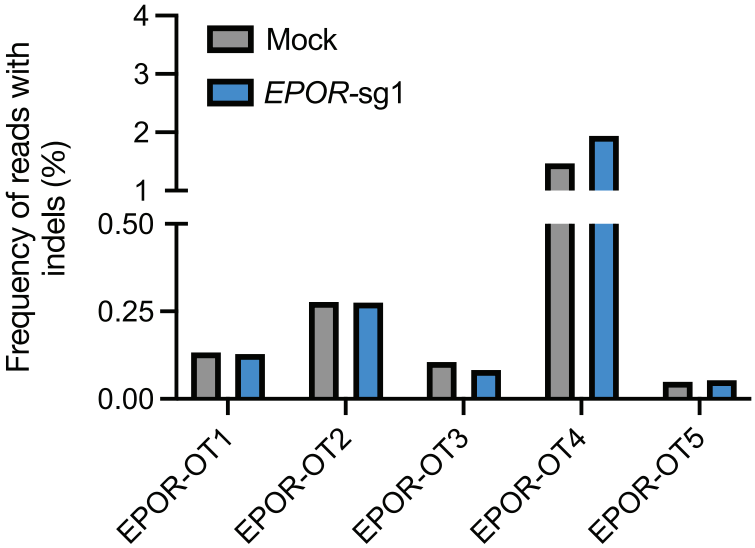

**Extended Data Figure 7. Off-target analysis of *EPOR*-sg1 by *in silico* prediction and targeted next-generation sequencing (NGS).**

**a.** Complete list of potential off-target sites predicted by COSMID with a score $\leq$5.5. **b.** Summary of off-target sites classified by region in genome. **c.** Detailed summary of 5 candidate off-target sites found in an exon or UTR of the genome. Sequence nucleotide mismatches highlighted in red, inserted nucleotides underlined, and missing nucleotides indicated by "^" symbol. PAM sequence highlighted in blue. **d.** Sequencing read depth of mock edited and *EPOR*-sg1 edited sample sent for each off-target site. N=1 biological HSPC donor. **e.** Frequency of indels detected in each sample at each off-target site. N=1 biological HSPC donor.

### Extended Data Figure 8

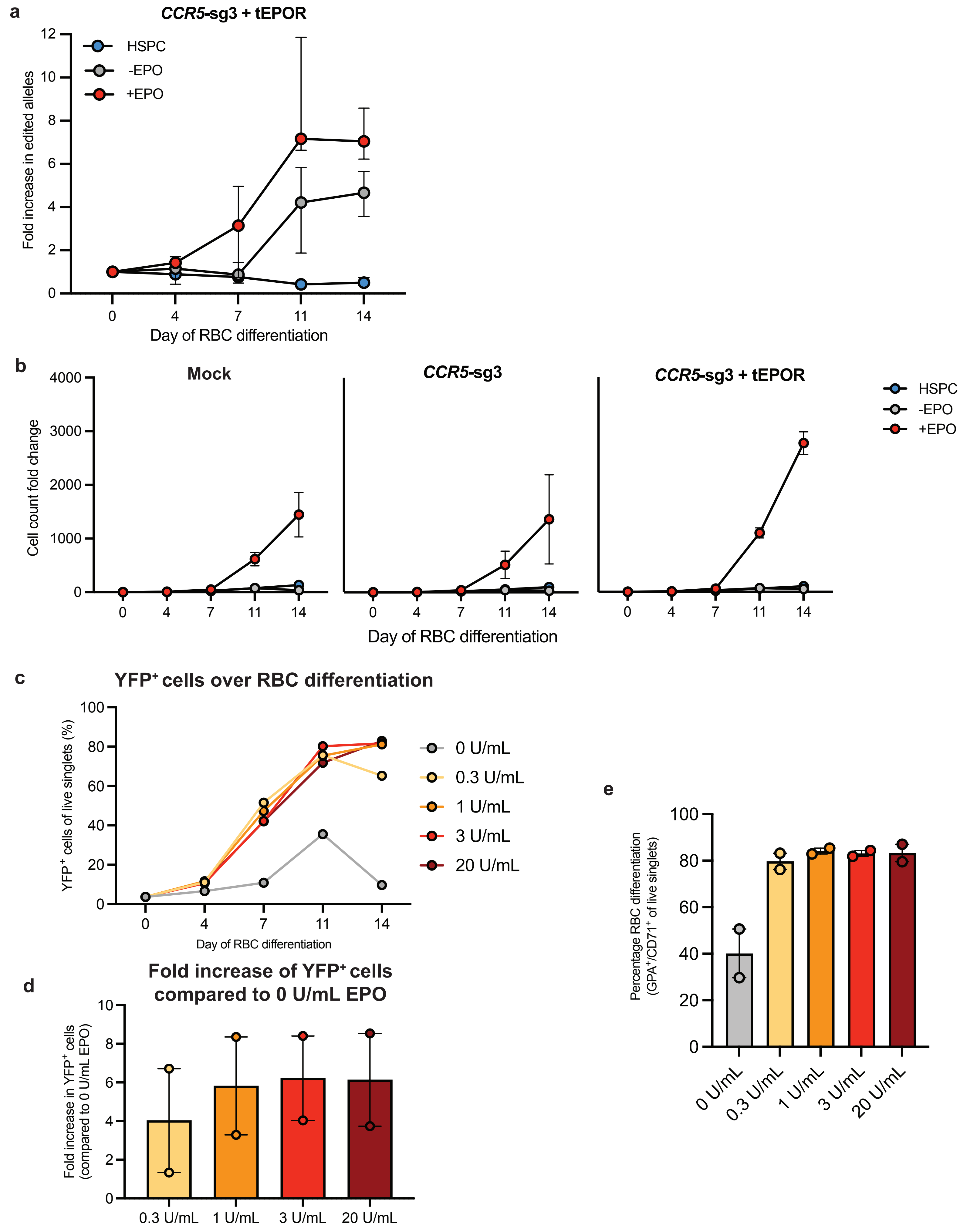

**Extended Data Figure 8. Safe harbor integration of *tEPOR* leads to increased erythroid proliferation and editing frequencies with little impact at different concentrations of EPO.**

**a.** Fold change in edited alleles at *CCR5* over course of RBC differentiation +/- EPO or maintained in HSPC media measured by ddPCR. Points shown as median ± interquartile range. N=3-4 biologically independent HSPC donors. **b.** Cell count fold change in mock and edited cells maintained in RBC media +/- EPO or HSPC media. Points represent mean ± SEM. Values represent biologically independent HSPC donors N=3 for Mock and *CCR5*-sg3 + *tEPOR* and N=2 for *CCR5*-sg3. **c.** Percentage of $YFP^+$ cells of live single cells throughout RBC differentiation with 0 U/mL – 20 U/mL EPO from one biological HSPC donor. **c.** Fold increase of $YFP^+$ cells at each concentration of EPO compared to 0 U/mL EPO at day 14 of differentiation. Bars represent median ± 95% confidence interval. N=2 biological HSPC donors. **d.** Percentage of $GPA^+/CD71^+$ of live single cells on day 14 of differentiation. Bars represent median ± 95% confidence interval. Values represent N=2 biologically independent HSPC donors.

### Extended Data Figure 9

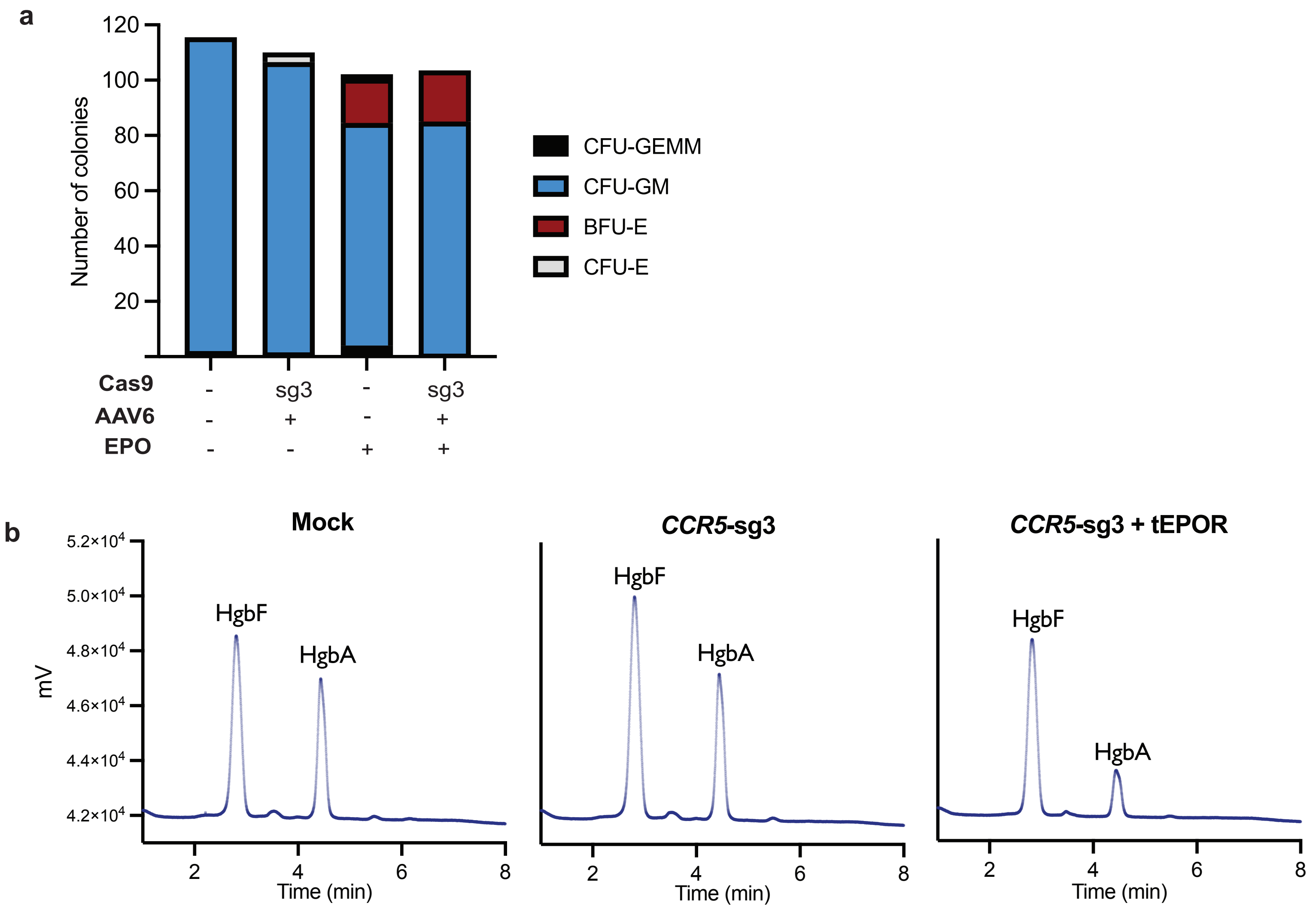

**Extended Data Figure 9. Safe harbor integration of *tEPOR* does not affect HSPC lineage or hemoglobin formation.**

**a.** CFU assay of mock edited cells versus cells edited with *CCR5*-sg3 + *tEPOR.* Bars represent total number of colonies of each type: CFU-GEMM (multi-potential granulocyte, erythroid, macrophage, megakaryocyte progenitor cells), CFU-GM (colony forming unit-granulocytes and monocytes), BFU-E (erythroid burst forming units), CFU-E (colony forming unit-erythroid) colonies. N=1. Mock condition from Extended Data 3c shown here for comparison. **b.** Representative HPLC plots of cells targeted with *CCR5*-sg3 or *CCR5*-sg3 + *tEPOR* at day 14 of RBC differentiation. HgbF=fetal hemoglobin, HgbA=adult hemoglobin.

#### Extended Data Figure 10

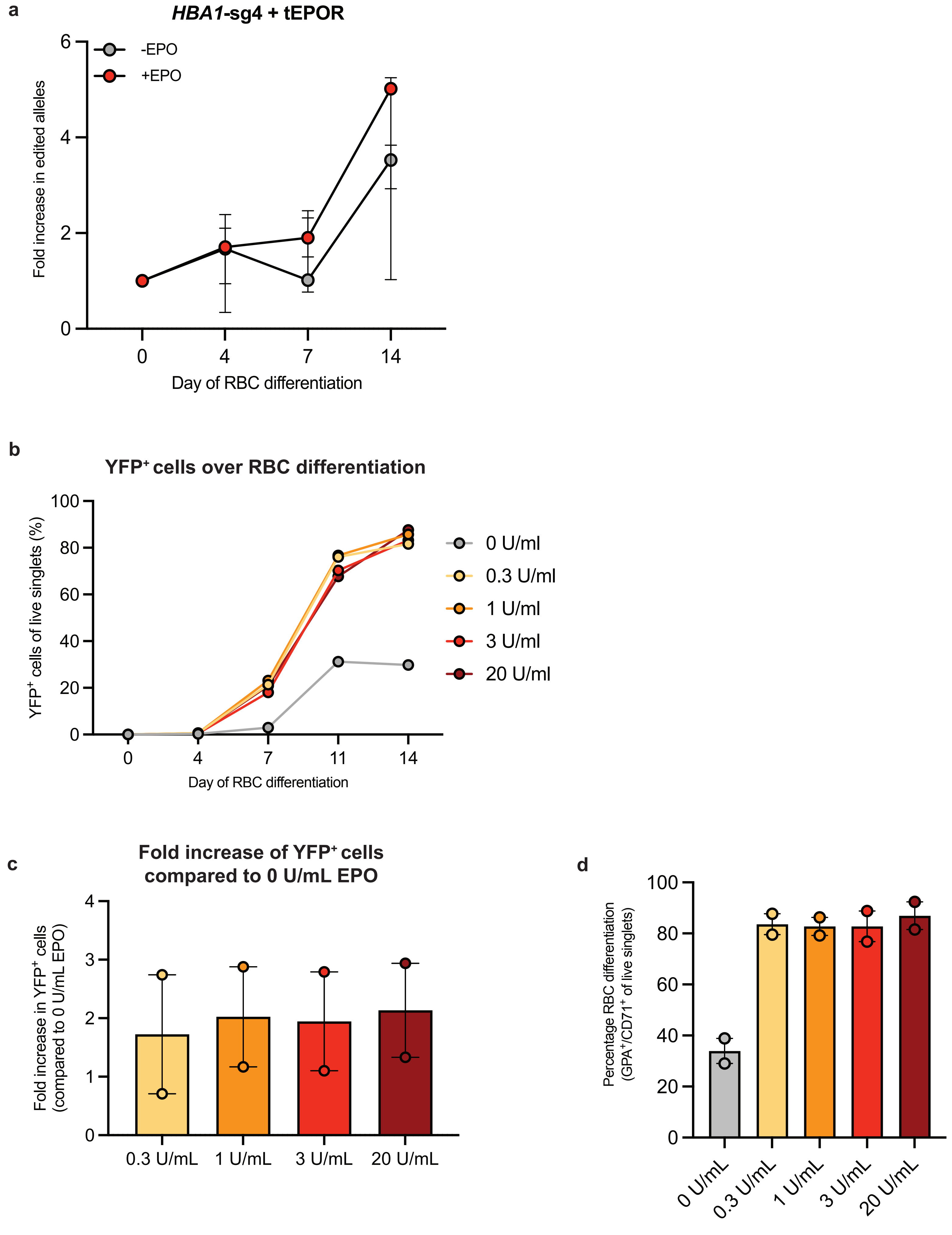

**Extended Data Figure 10. Erythroid specific expression of *tEPOR* leads to increased editing frequencies and is not affected by EPO concentration.**

**a.** Fold change in edited alleles at *HBA1* over course of RBC differentiation +/- EPO measured by ddPCR. Points shown as median ± 95% CI. N=3 biologically independent HSPC donors. **b.** Percentage of $YFP^+$ cells of live single cells throughout RBC differentiation with 0 U/mL – 20 U/mL EPO in one biological HSPC donor. **c.** Fold increase of $YFP^+$ cells at each EPO concentration compared to 0 U/mL EPO at day 14 of RBC differentiation. Bars represent median ± 95% confidence interval. N=2 biological HSPC donors. **d.** Percentage of $GPA^+/CD71^+$ of live single cells on day 14 of differentiation. Bars represent median ± 95% confidence interval. Values represent N=2 biologically independent HSPC donors.

#### Extended Data Figure 11

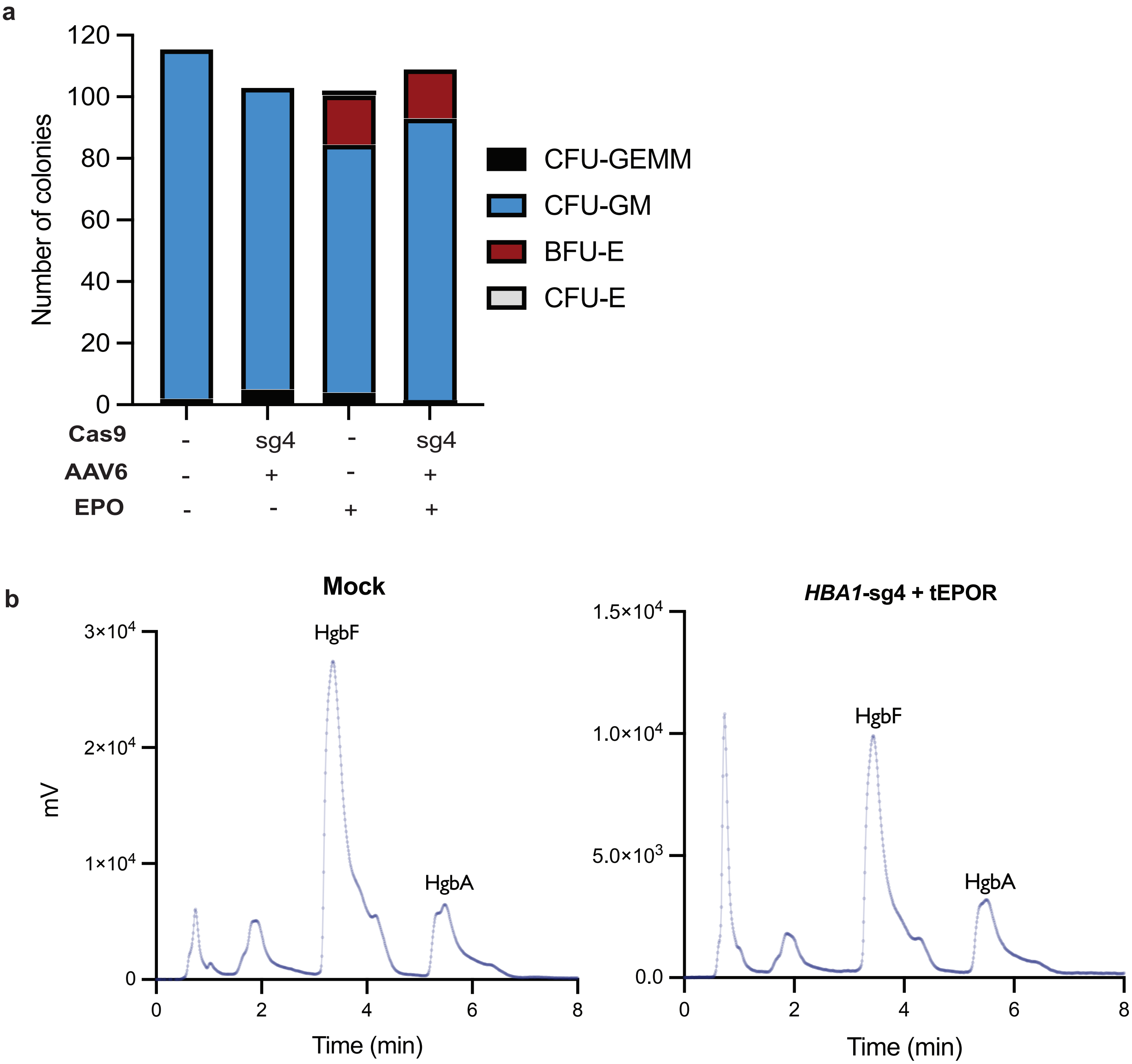

**Extended Data Figure 11. Erythroid specific expression of *tEPOR* does not affect HSPC lineage or hemoglobin formation.**

**a.** CFU assay of mock edited cells versus cells edited with *HBA1*-sg4 + *tEPOR.* Bars represent total number of colonies of each type: CFU-GEMM (multi-potential granulocyte, erythroid, macrophage, megakaryocyte progenitor cells), CFU-GM (colony forming unit-granulocytes and monocytes), BFU-E (erythroid burst forming units), CFU-E (colony forming unit-erythroid) colonies. N=1. Mock condition from Extended Data 3c shown here for comparison. **b.** Representative HPLC plots of cells targeted with *HBA1*-sg4 + *tEPOR* at day 14 of RBC differentiation. HgbF=fetal hemoglobin, HgbA=adult hemoglobin.

#### Extended Data Figure 12

**a**

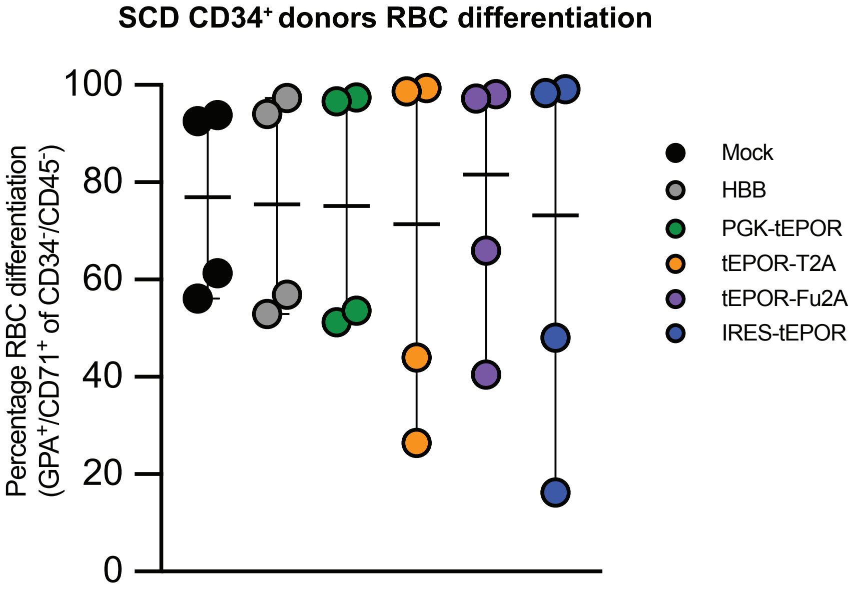


**b**

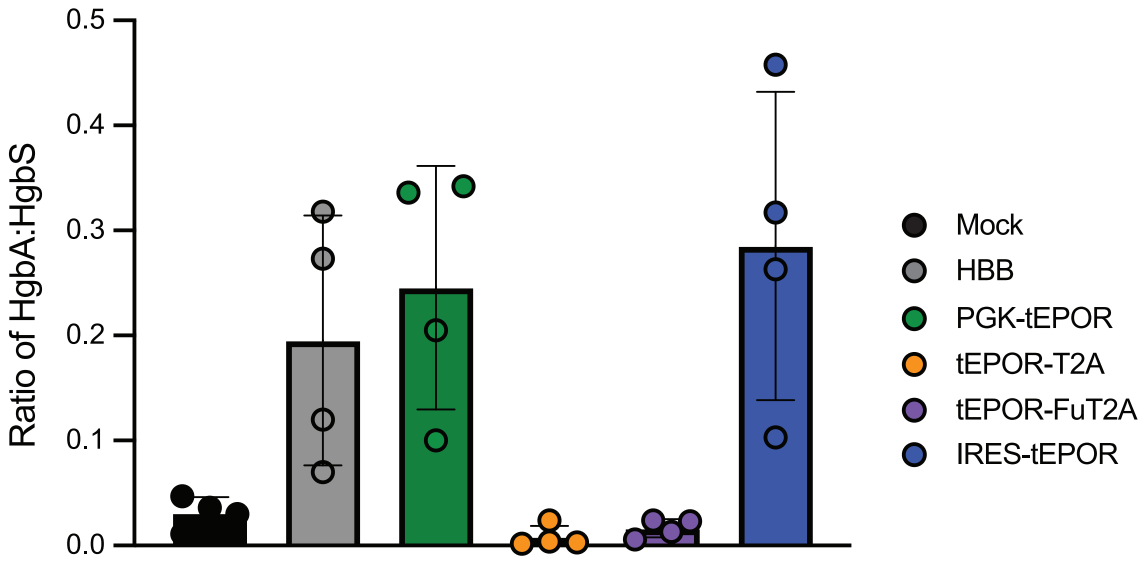

**Extended Data Figure 12. Hemoglobin tetramer analysis of *tEPOR* edited SCD patient HSPCs following RBC differentiation.**

**a.** Percentage of $GPA^{+}/CD71^{+}$ of $CD34^{-}/CD45^{-}$ cells on day 14 of RBC differentiation as determined by flow cytometry. Points shown as median ± 95% confidence interval. N=4 biologically independent HSPC donors. **b.** Ratio of adult hemoglobin to sickle hemoglobin from HPLC analysis of hemoglobin tetramers from differentiated SCD patient HSPCs. Bars shown as mean ± SD. N=4 biologically independent HSPC donors.

#### Extended Data Figure 13

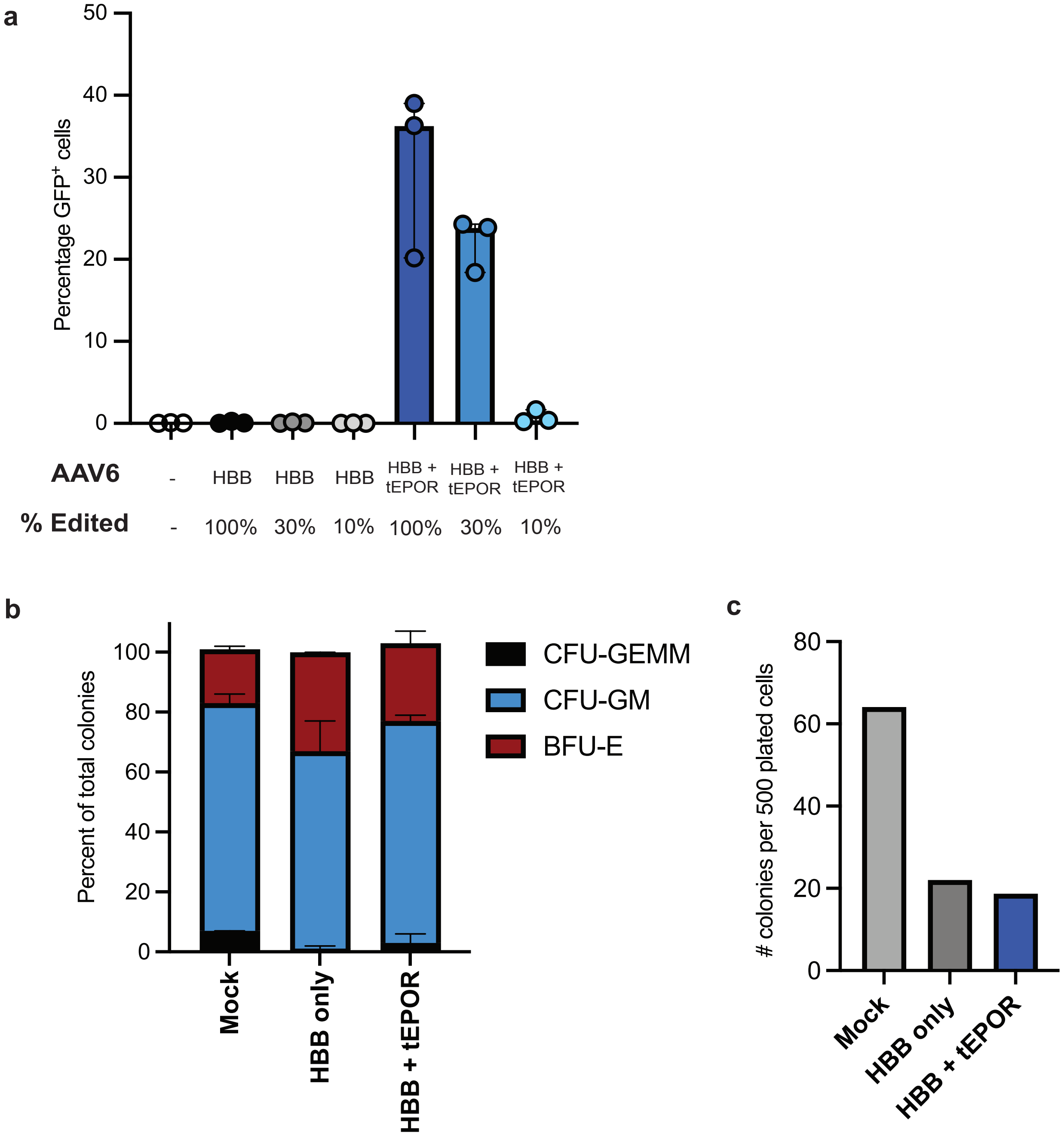

**Extended Data Figure 13. Multiplexed edited strategy shows maintenance of *EPOR* truncation at day 14 of RBC differentiation and does not disrupt HSPC lineage formation.**

**a.** Percentage of GFP$^+$ cells of live single cells on day 14 of RBC differentiation as determined by flow cytometry. Points represent median ± interquartile range. Values represent N=3 biologically independent HSPC donors. **b.** CFU assay of mock, single edited, and multiplex edited HSPCs. Bars represent percent of total colonies: CFU-GEMM (multi-potential granulocyte, erythroid, macrophage, megakaryocyte progenitor cells), CFU-GM (colony forming unit-granulocytes and monocytes), and BFU-E (erythroid burst forming units). N=1. **c.** Number of total colonies in CFU assay produced per 500 plated cells in each condition.

#### Supplementary Figure 1

**a**

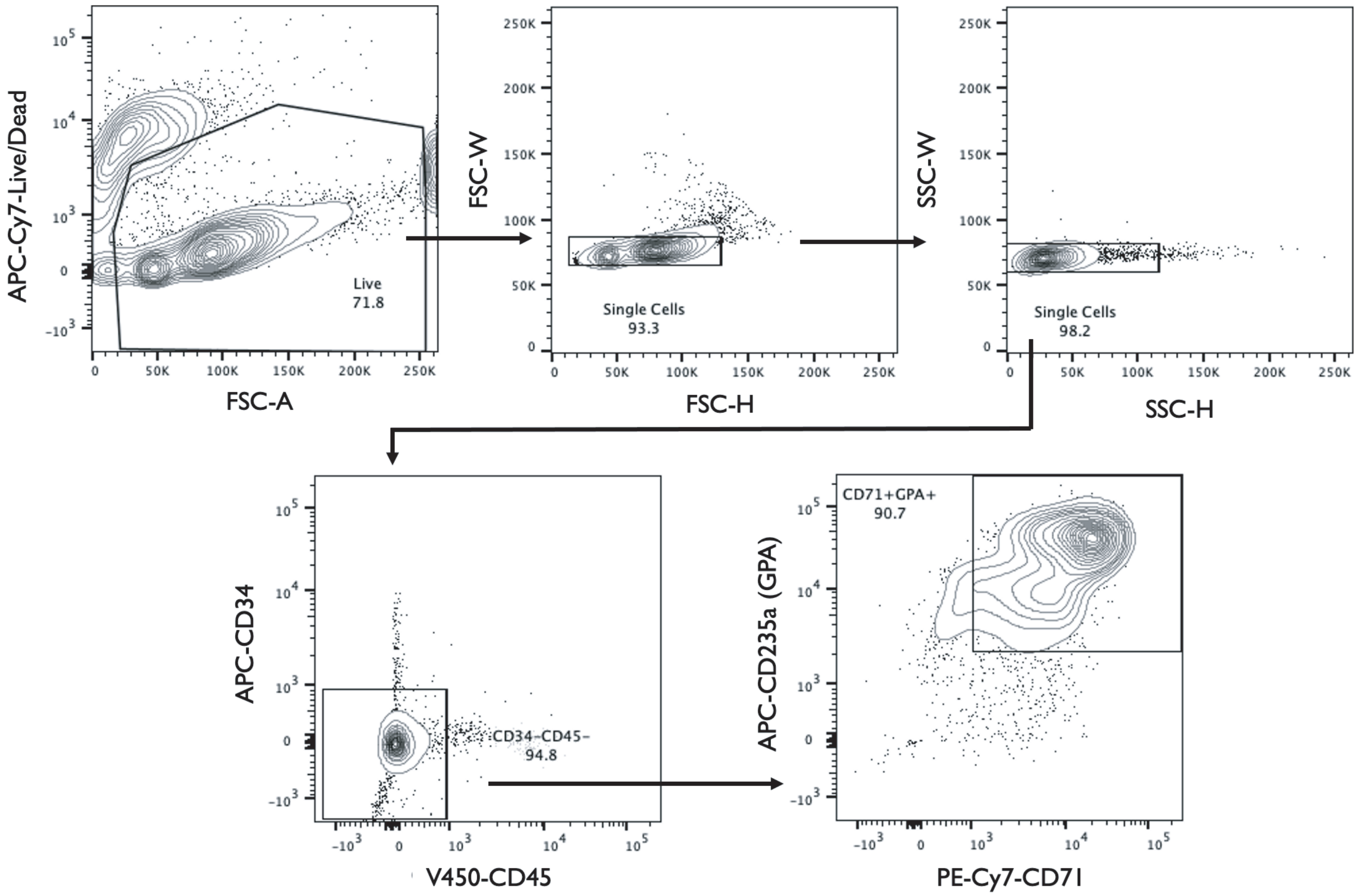

**Supplementary Figure 1. Gating strategy for flow cytometry of RBC differentiation**

**a.** Representative flow plots of the gating strategy used to quantify RBC differentiation.
